# Correction of the scientific production: publisher performance evaluation using a dataset of 4844 PubMed retractions

**DOI:** 10.1101/2022.01.23.477400

**Authors:** Catalin Toma, Liliana Padureanu, Bogdan Toma

## Abstract

**Background:** Withdrawal of problematic scientific articles after publication is one of the mechanisms for correcting the literature available to publishers, especially in the conditions of the ever-increasing trend of publishing activity in the medical field. The market volume and the business model justify publishers’ involvement in the post-publication quality control(QC) of scientific production. The limited information about this subject determined us to analyze retractions and the main retraction reasons for publishers with many withdrawn articles. We also propose a score to measure the evolution of their performance. The data set used for this article consists of 4844 PubMed retracted papers published between 1.01.2009 and 31.12.2020.

**Methods:** We have analyzed the retraction notes and retraction reasons, grouping them by publisher. To evaluate performance, we formulated an SDTP score whose calculation formula includes several parameters: speed (article exposure time(ET)), detection rate (percentage of articles whose retraction is initiated by the editor/publisher/institution without the authors’ participation), transparency (percentage of retracted articles available online and clarity of retraction notes), precision (mention of authors’ responsibility and percentage of retractions for reasons other than editorial errors).

**Results:** The 4844 withdrawn articles were published in 1767 journals by 366 publishers, the average number of withdrawn articles/journal being 2.74. Forty-five publishers have more than ten withdrawn articles, holding 88% of all papers and 79% of journals. Combining our data with data from another study shows that less than 7% of PubMed journals withdrew at least one article. Only 10.5% of the withdrawal notes included the individual responsibility of the authors. Nine of the top 11 publishers had the largest number of articles withdrawn in 2020, in the first 11 places finding, as expected, some big publishers. Retraction reasons analysis shows considerable differences between publishers concerning the articles ET: median values between 9 and 43 months (mistakes), 9 and 73 months (images), 10 and 42 months (plagiarism & overlap).

The SDTP score shows, between 2018 and 2020, an improvement in QC of four publishers in the top 11 and a decrease in the gap between 1st and 11th place. The group of the other 355 publishers also has a positive evolution of the SDTP score.

**Conclusions:** Publishers have to get involved actively and measurably in the post-publication evaluation of scientific products. The introduction of reporting standards for retraction notes and replicable indicators for quantifying publishing QC can help increase the overall quality of scientific literature.

## Introduction

> *„One of the greatest criticisms in the blogosphere is not so much that the current rules and guidelines are weak or poor, but that enforcement and irregular application of those rules, particularly by COPE member journals and publishers, confuses the readership, disenfranchises authors who remain confused(despite having a stricter and more regulated system) and provides an imbalanced publishing structure that has weak, or limited, accountability or transparency.”*(Teixeira da Silva and Dobránszki 2017)

The publication of scientific literature represents, globally, a market of considerable size, which reached a record value of $ 28 billion in 2019 (from 9,4 billion in 2011 (van Noorden 2013)), fell to $ 26.5 billion in 2020, with forecasts suggesting a recovery of losses by 2023. Revenues from the publication of articles in 2019 were $ 10,81 billion, those from the publication of books $ 3,19 billion, derivative products represent the difference. The segment of medical publications (12.8 billion in 2020) is constantly growing. Estimations show that in 2024 the medical literature will exceed the volume of technical and scientific literature(International Association of Scientific, Technical and Medical Publishers 2021). The continued growth was accompanied by a consolidation process that made the top 5 publishers in 2013 represent over 50% of all published articles. These changes occur in an atypical market where publishers have high-profit margins (Hagve 2020), do not pay for purchased goods (authors are not paid), do not pay for quality control (peer-review), and have a monopoly on the content of published articles (Ingelfinger law)((Larivière et al. 2015).

Under these conditions, improving the quality of published scientific production should be a priority for publishers. One of the methods is the withdrawal of invalid articles from a scientific, ethical or legal point of view (questionable research practices-QRP, questionable publication practices-QPP). The interest in correcting the literature seems to be confirmed by recent developments: the number of journals with at least one withdrawn article increased from 44 in 1997 to 488 in 2016. (Brainard 2018). The continued growth may be due to improved capacity to detect and remove problematic articles (Vuong et al. 2020), which, despite a somewhat reluctant if not resisting editorial environment (Marcus and Oransky 2014; Friedman et al. 2020) and the lack of significant progress in reporting (Teixeira da Silva and Dobránszki 2017; Vuong 2020) continues to expand in the publishing environment. For example, in the case of PubMed retractions, the year 2020 was a record year in terms of withdrawal notes, targeting 878 articles published in more than 12 years. (Toma and Padureanu 2021)

The withdrawals in the biomedical journals indexed in PubMed represent an intensely researched topic in the last two decades, numerous articles making valuable contributions in this field (Nath et al. 2006; Redman et al. 2008; Wager and Williams 2011; Steen 2011; Samp et al. 2012; Fang et al. 2012; Steen et al. 2013; Decullier et al. 2013; Madlock-Brown and Eichmann 2015; Mongeon and Larivière 2016; Rosenkrantz 2016; Pantziarka and Meheus 2019; Rapani et al. 2020; Bhatt 2021). However, there is little information on the article retractions at the publisher level and, therefore, an incomplete picture of the challenges/difficulties they face in the post-publication quality control of products delivered to consumers of scientific information. Several authors point out issues that arise when withdrawing a scientific article:

- The process of withdrawing an article is a complex one, depending on several factors: who initiates the withdrawal, the context, the communication between the parties, the editorial experience (Williams and Wager 2013).
- The clarity of the withdrawal notes leaves much to be desired and presents a significant variability between journals and/or publishers in relation to the COPE guidelines (Bilbrey et al. 2014; Cox et al. 2018; Coudert 2019; Teixeira da Silva and Dobránszki 2017), although the need for a uniform approach has long been required (Fang et al. 2012).
- The individual contribution of the authors is rarely mentioned in the withdrawal notes, contrary to the COPE recommendations (Coudert 2019).
- The online presence of withdrawn articles, required in the COPE guidelines, ensures more transparency and avoids the occurrence of “silent or stealth retractions” (Teixeira da Silva 2016).
- The role of publishers (avoiding what is called editorial misconduct) in the process of correcting the scientific literature is an important one(Shelomi 2014). Editorial errors (duplicate publication, accidental publication of wrong version / rejected article, wrong journal publication) were identified in different proportions: 7,3% (328 cases) in a 2012 study that analyzed 4449 articles withdrawn between 1928-2011 (Grieneisen and Zhang 2012), 1,5% (5 cases) in another study (Coudert 2019), 5% in the Bar-Ilan study (Bar-Ilan and Halevi 2018), 3,7% in our study on PubMed withdrawals between 2009 and 2020 (Toma and Padureanu 2021).
- The level of involvement of publishers and editors in article withdrawal is variable (Grieneisen and Zhang 2012; Cox et al. 2018), although most of them can initiate the withdrawal of an article without the authors’ consent (Resnik et al. 2015).
- Different efficiency of QRP and QPP detection mechanisms at the editorial level may explain the differences between publishers (Fanelli 2013), the possible application of postpublication peer review (Teixeira da Silva and Dobránszki 2015; Ali and Watson 2016) being able to contribute both to the increase of the detection capacity and the reduction of the differences between journals/publishers.

As many editorial policies are / can be implemented at the publisher level and positively / negatively affect the performance of all journals in its portfolio, we thought it would be helpful to present an overview of withdrawn articles and a structure of reasons for withdrawal for major publishers using a dataset obtained from the analysis of 4844 biomedical articles indexed in PubMed and withdrawn between 2009-2020.

We also consider it worthwhile to initiate a debate on the performance of publishers in correcting the scientific literature. For this reason, we propose a score based on four indicators: speed of article withdrawal, post-publication ability of the publisher/editors to detect QRP/QPP articles, the transparency of withdrawals (measured by the online maintenance of withdrawn articles and the clarity of withdrawal notes) and the precision of the correction process (identification of those responsible and the degree of avoidance of editorial errors).

What do we think this article brings new?

- Presentation of the main retraction reasons and exposure time (ET) for leading publishers;
- Dynamics of withdrawal notes at publisher level;
- Evaluation of publisher performance for three main retraction reasons;
- Proposal of a tool for measuring publisher post-publication QC performance (SDTP score: Speed-Detection-Transparency-Precision score)

Limitations.

- The databases used showed some errors that can lead to changes in the exposure time, although in a minimal number.
- The interpretation of retraction notes may generate classification errors, when several retraction reasons are mentioned;
- Modifications/completions made after the study by publishers or editors to the registrations on their sites may modify the figures obtained by us;
- The score is obtained by simple summation without taking into account the lower / higher weight that can be assigned to a specific component.

## Materials and methods

The methodology used to collect the data is presented in detail in another article (Toma and Padureanu 2021).

For the publishing and editorial performance indicators, we have built a score(SDTP score) consisting of 4 components and six values represented equally and calculated from the data set collected: speed, detection rate, transparency, and precision.

In case of publisher/editor involvement(2-Detection rate), we used in the calculation all articles that mentioned in the withdrawal note involvement of publisher, editor in chief, editorial board, institution, Office of Research Investigation, without authors.

In order to measure the identification of the authors’ responsibility (5-Precision-Individual responsibility), we used as calculation base the number of articles with more than one author and retractions for other reasons than editorial errors.

### Calculation of SDTP score

The values for each publisher are compared with the values of the entire set of 4844 retracted articles(2009-2020), 3931 articles(2009-2019), and 3361 articles(2009-2018).

There are two situations:

A. Values above average are considered poor performance. Example: exposure time(speed).

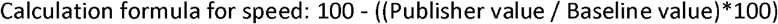
B. A percentage value above the average is considered good performance. Example: detection rate, percentage of online papers, percentage of clear retraction notes, percentage of retractions in which individual author responsibilities are mentioned, percentage of retractions not due to editorial errors.

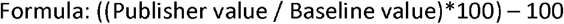

The values obtained are summed and form the SDTP score of the publisher.

## Results

### Retractions by Publishers

The 4844 retracted articles were published in 1767 journals. The average number of withdrawn articles is 2,74/journal.

Several studies have reported publisher rankings, with the top positions being consistently occupied by publishers with a large number of publications (Cox et al. 2018; Tripathi et al. 2019; Vuong et al. 2020). This is also reflected in the results obtained by us. Forty five publishers with more than 10 retractions account for 88% (n = 4261) of retractions and 79% (n = 1401) of journals(table 2). The remaining 583 papers are published in 366 journals associated with 321 publishers.

**Table 1.**
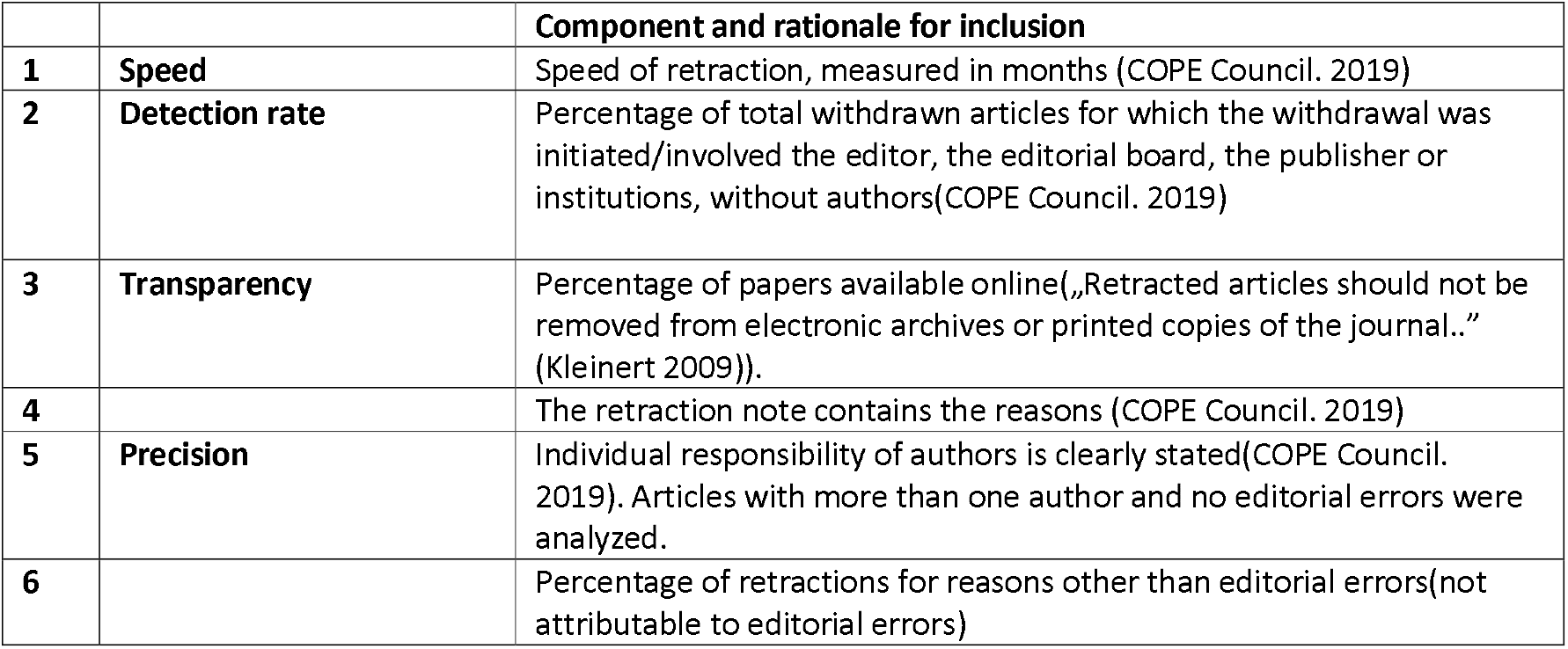
SDTP score components

|  |  | Component and rationale for inclusion |
| --- | --- | --- |
| 1 | Speed | Speed of retraction, measured in months (COPE Council. 2019) |
| 2 | Detection rate | Percentage of total withdrawn articles for which the withdrawal was initiated/involved the editor, the editorial board, the publisher or institutions, without authors (COPE Council. 2019) |
| 3 | Transparency | Percentage of papers available online („Retracted articles should not be removed from electronic archives or printed copies of the journal..” (Kleinert 2009)). |
| 4 |  | The retraction note contains the reasons (COPE Council. 2019) |
| 5 | Precision | Individual responsibility of authors is clearly stated (COPE Council. 2019). Articles with more than one author and no editorial errors were analyzed. |
| 6 |  | Percentage of retractions for reasons other than editorial errors (not attributable to editorial errors) |

**Table 2.** Publishers with more than ten retracted articles.

|  | Publisher | Articles | Journals | /journal |
| --- | --- | --- | --- | --- |
| 1 | ELSEVIER | 846 | 309 | 2,74 |
| 2 | SPRINGER NATURE | 749 | 305 | 2,46 |
| 3 | WOLTERS KLUWER | 346 | 142 | 2,45 |
| 4 | WILEY-BLACKWELL | 325 | 171 | 1,90 |
| 5 | PLOS | 275 | 7 | 39,28 |
| 6 | SAGE | 248 | 47 | 5,28 |
| 7 | TAYLOR AND FRANCIS | 223 | 95 | 2,35 |
| 8 | HINDAWI | 140 | 52 | 2,69 |
| 9 | DOVE MEDICAL PRESS | 111 | 23 | 4,83 |
| 10 | E-CENTURY PUBLISHING | 71 | 7 | 10,14 |
| 11 | SPANDIDOS PUBLICATIONS | 71 | 7 | 10,14 |
| 12 | OXFORD UNIVERSITY PRESS | 65 | 39 | 1,67 |
| 13 | MARY ANN LIEBERT | 63 | 26 | 2,42 |
| 14 | VERDUCI EDITORE | 56 | 1 | 56 |
| 15 | AMERICAN CHEMICAL SOCIETY | 53 | 17 | 3,12 |
| 16 | AMERICAN ASSOCIATION FOR CANCER RESEARCH | 49 | 6 | 8,17 |
| 17 | AMERICAN SOCIETY FOR MICROBIOLOGY | 44 | 10 | 4,40 |
| 18 | FRONTIERS MEDIA | 44 | 18 | 2,44 |
| 19 | MULTIDISCIPLINARY DIGITAL PUBLISHING INSTITUTE | 42 | 17 | 2,47 |
| 20 | NATIONAL ACADEMY OF SCIENCES | 36 | 1 | 36 |
| 21 | PORTLAND PRESS | 32 | 3 | 10,67 |
| 22 | BMJ PUBLISHING GROUP | 29 | 17 | 1,70 |
| 23 | AMERICAN PHYSIOLOGICAL SOCIETY | 26 | 8 | 3,25 |
| 24 | FUNDACAO DE PESQUISAS CIENTIFICAS DE RIBEIRAO PRETO | 20 | 1 | 20 |
| 25 | AMERICAN MEDICAL ASSOCIATION | 19 | 12 | 1,58 |
| 26 | SOCIETY FOR NEUROSCIENCE | 18 | 1 | 18 |
| 27 | AMERICAN ASSOCIATION FOR THE ADVANCEMENT OF SCIENCE | 17 | 3 | 5,66 |
| 28 | AMERICAN SOCIETY OF HEMATOLOGY | 17 | 1 | 17 |
| 29 | AMERICAN SOCIETY FOR PHARMACOLOGY AND EXPERIMENTAL THERAPEUTICS | 16 | 3 | 5,33 |
| 30 | BENTHAM | 16 | 8 | 2 |
| 31 | CELL PHYSIOL BIOCHEM PRESS | 16 | 1 | 16 |
| 32 | IMPACT JOURNALS | 16 | 2 | 8 |
| 33 | MEDICAL SCIENCE INTERNATIONAL PUBLISHING | 16 | 1 | 16 |
| 34 | AMERICAN DIABETES ASSOCIATION | 15 | 2 | 7,5 |
| 35 | AMERICAN SOCIETY OF CLINICAL INVESTIGATION | 15 | 1 | 15 |
| 36 | CUREUS, INC. | 14 | 1 | 14 |
| 37 | THE COMPANY OF BIOLOGISTS LTD. | 13 | 2 | 6,5 |
| 38 | KOWSAR PUBLISHING COMPANY | 12 | 3 | 4 |
| 39 | KARGER | 12 | 8 | 1,5 |
| 40 | AME PUBLISHING COMPANY | 11 | 4 | 2,75 |
| 41 | AMERICAN THORACIC SOCIETY | 11 | 3 | 3,66 |
| 42 | ASSOCIACAO BRASILEIRA DE DIVULGACAO CIENTIFICA | 11 | 1 | 11 |
| 43 | FUTURE MEDICINE LTD | 11 | 9 | 1,22 |
| 44 | INTERNATIONAL INSTITUTE OF ANTICANCER RESEARCH | 11 | 1 | 11 |
| 45 | IOS Press | 10 | 5 | 2 |

**Table 3.**
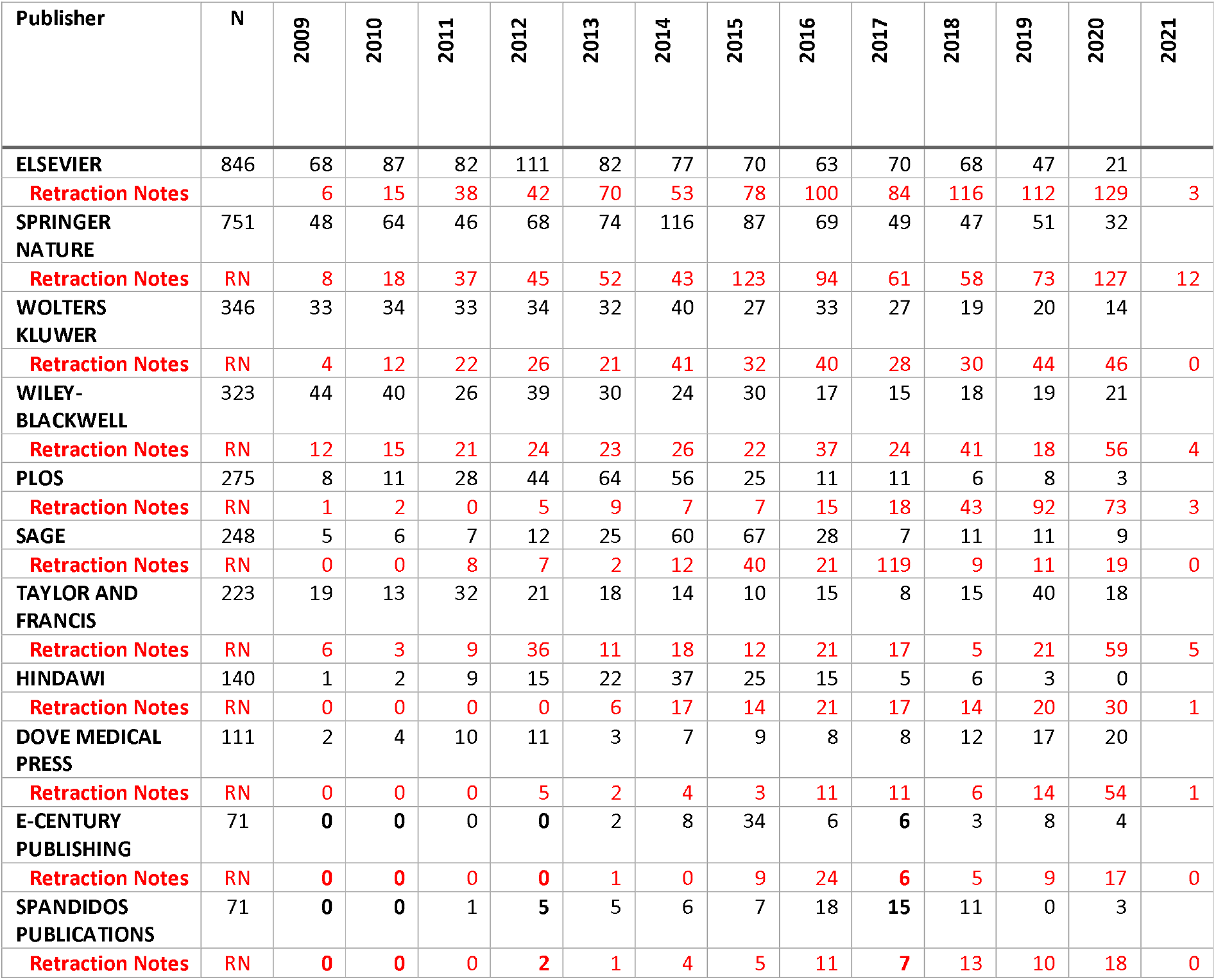
Retracted articles and retraction notes by year for top 11 publishers.

The top 11 publishers have 3,405 withdrawn articles(70.3%) in 1165 journals (65.9%). In the following, we will only analyze their evolution and performance. The rest of the publishers will be analysed within a single group.

### Retraction notes/publisher(2009-2020)

2020 is the most consistent year for retracted articles for almost all publishers in the top 11, except PLOS, which peaked in 2019, SAGE in 2017, and E-Century Publishing in 2015. The period 2012-2014 seems to be, for most publishers, the beginning of a greater interest in correcting the medical literature.

### Publishers and retraction reasons

Retraction reasons for the top 11 publishers are presented in table 4. Multiple reasons in one retraction note were added to the respective categories, thus explaining the publisher percentages sum higher than 100%.

**Table 4.** Retraction reasons for top 11 publishers(3405 retracted papers, 70,3%).

| *Publisher* |  | *MISTAKES* | *IMAGES* | *PLAGIARISM* | *OVERLAP* | *FRAUD* | *ETHICS* | *AUTHORSHIP* | *UNCLEAR* | *EDITOR* | *PROPERTY* | *OTHER* |
| --- | --- | --- | --- | --- | --- | --- | --- | --- | --- | --- | --- | --- |
| *ELSEVIER* | **846** | 32,98% | **33,2%** | 9,8% | 10,87% | 2,6% | 6,38% | 4,73% | 5,79% | 4,25% | 1,3% | 1,3% |
|  | **N** | **279** | **281** | **83** | **92** | **22** | **54** | **40** | **49** | **36** | **11** | **11** |
| *SPRINGER NATURE* | **749** | 29,64% | 17,62% | 18,42% | 12,02% | 17,09% | 8,94% | 5,74% | 0,5% | 2,4% | 4,67% | 1,07% |
|  | **N** | **222** | **132** | **138** | **90** | **128** | **67** | **43** | **4** | **18** | **35** | **8** |
| *WOLTERS KLUWER* | **346** | 28,61% | 6,94% | 24,85% | 15,6% | 1,73% | 8,67% | 6,36% | 6,65% | 6,94% | 1,73% | 1,73% |
|  | **N** | **99** | **24** | **86** | **54** | **6** | **30** | **22** | **23** | **24** | **6** | **6** |
| *WILEY-BLACKWELL* | **325** | 36% | 18,8% | 9,2% | 14,5% | 1,5% | 9,5% | 7,1% | 2,2% | 6,5% | 2,2% | 1,8% |
|  | **N** | **117** | **61** | **30** | **47** | **5** | **31** | **23** | **7** | **21** | **7** | **6** |
| *PLOS* | **275** | 29,5% | **63,3%** | 4% | 4% | 1,1% | 16,4% | 5,1% | -- | 0,7% | 1,1% | 0,4% |
|  | **N** | **81** | **174** | **11** | **11** | **3** | **45** | **14** | **--** | **2** | **3** | **1** |
| *SAGE* | **248** | 16,9% | 0,8% | 9,3% | 8,1% | **62,9%** | 4,8% | 7,3% | 1,2% | 2,4% | 1,2% | -- |
|  | **N** | **42** | **2** | **23** | **20** | **156** | **12** | **18** | **3** | **6** | **3** | **--** |
| *TAYLOR AND FRANCIS* | **223** | 26,9% | 11,7% | 13% | 17% | 13% | 6,7% | 10,3% | 1,8% | 6,7% | 2,7% | 1,3% |
|  | **N** | **60** | **26** | **29** | **38** | **29** | **15** | **23** | **4** | **15** | **6** | **3** |
| *HINDAWI* | **140** | 22,1% | 27,1% | **33,6%** | 16,4% | 0,7% | 9,3% | 10% | 0,7% | -- | 2,1% | -- |
|  | **N** | **31** | **38** | **47** | **23** | **1** | **13** | **14** | **1** | **--** | **3** | **--** |
| *DOVE MEDICAL PRESS* | **111** | 27,9% | **39,6%** | 14,4% | 16,2% | 2,7% | 14,4% | 4,5% | -- | -- | 1,8% | -- |
|  | **N** | **31** | **44** | **16** | **18** | **3** | **16** | **5** | **--** | **--** | **2** | **--** |
| *E-CENTURY PUBLISHING* | **71** | 26,8% | 2,8% | **36,6%** | 1,4% | 8,5% | 2,8% | 4,2% | -- | 25,4% | 1,4% | -- |
|  | **N** | **19** | **2** | **26** | **1** | **6** | **2** | **3** | **--** | **18** | **1** | **--** |
| *SPANDIDOS PUBLICATIONS* | **71** | **33,8%** | 29,6% | 28,2% | 5,6% | 1,4% | 7% | 11,3% | -- | -- | 1,4% | 1,4% |
|  | **N** | **24** | **21** | **20** | **4** | **1** | **5** | **8** | **--** | **--** | **1** | **1** |

**Table 5.** Mistakes/Inconsistent data per publisher

|  | Articles | ET(95% CI) | Median | Range | IQR |
| --- | --- | --- | --- | --- | --- |
| **Elsevier** | 279 | **29,03**(25,87-32,19) | 20 | 133 | 33 |
| **Springer Nature** | 222 | **27,76**(24,04-31,48) | 17 | 140 | 29 |
| **Wolters Kluwer** | 99 | **19,69**(15,26-24,12) | 11 | 99 | 16 |
| **Wiley-Blackwell** | 117 | **26,19**(22,45-29,93) | 23 | 98 | 26 |
| **PLOS** | 81 | **41,58**(34,92-48,24) | 43 | 117 | 57 |
| **SAGE** | 42 | **21,36**(16,05-26,67) | 17 | 61 | 27 |
| **Taylor & Francis** | 60 | **15,43**(11,34-19,53) | 12 | 72 | 16 |
| **Hindawi** | 31 | **29,74**(20,24-39,25) | 16 | 81 | 41 |
| **Dove Medical Press** | 31 | **31,90**(16,62-47,19) | 9 | 139 | 46 |
| **E-Century Publishing** | 19 | **22,58**(15,48-29,68) | 26 | 48 | 26 |
| **Spandidos Publications** | 24 | **17,17**(9,68-24,65) | 9 | 70 | 20 |

**Table 6.**
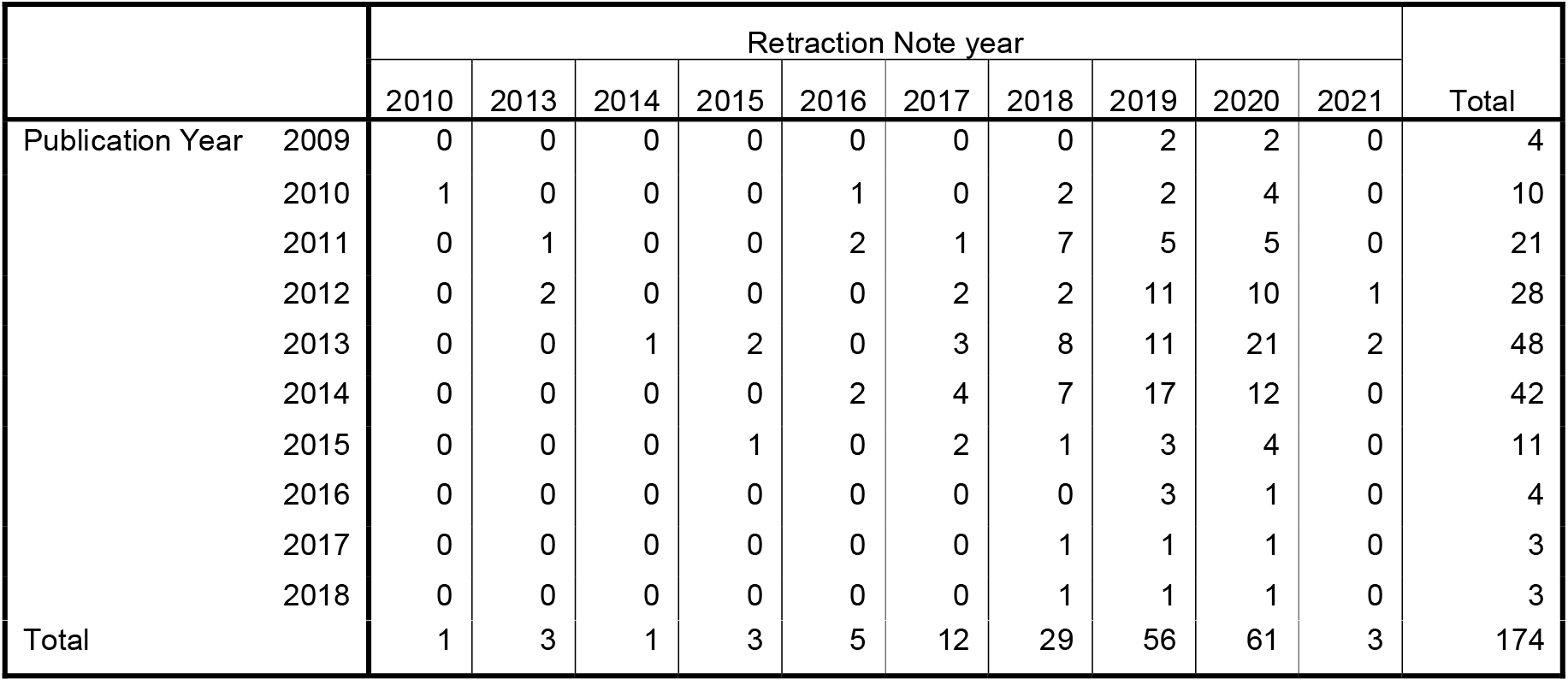
Evolution of PLOS image retractions.

#### Mistakes/Inconsistent data

Of the 1553 cases, 1005 (64.7%) belong to the top 11 publishers. The rest of the publishers account for 548/1553 cases. In 229/1553 cases (14.7%, 95% CI 12.9-16.5), the reason for the withdrawal was data fabrication. For the top 11 publishers, there are 127/1005 (12.6%, CI95% 12.6-14.7) cases of data fabrication (48 Elsevier, 22 for Springer Nature and Wolters Kluwer, 12 for Wiley-Blackwell, 7 for SAGE, 5 for E-Century Publishing, 4 for Taylor & Francis and PLOS, 3 for Hindawi). The other publishers have 102/548 articles retracted for data fabrication (18.6%, CI95% 15.3-21.8).

The lowest ET average belongs to Taylor & Francis(15,4 months) and the highest to PLOS(41,5 months). We note in the meantime, in most of the cases, skewed distributions and median values ranging from an encouraging nine months(Dove Medical Press and Spandidos Publications), 11 months(Wolters Kluwer), 12 months(Taylor&Francis) to a rather unexpected 43 months for PLOS.

#### Images

Several publishers have a high number of retractions due to image problems: PLOS(174 of 275 papers, 63,3%), Elsevier(281 of 846 papers, 33,2%), Springer Nature(132 of 749 papers, 17,62%), possibly signaling the implementation of procedures and technologies to detect problematic articles.

In the case of PLOS, out of the 174 articles withdrawn for image problems, 150 (86.2%) were published between 2011-2015, and 90.9% (158) of the withdrawal notes were published between 2017-2020. This suggests that 2017 could be the year when the systematic and retroactive verification of the images in the articles published in 2009-2020 began. Only ten articles published by PLOS between 2016-2020 (no articles in 2019 and 2020) are withdrawn due to images, suggesting the effectiveness of the measures implemented by this publisher and that probably, the articles with questionable images are stopped before publication.

In the case of Elsevier, out of the total of 281 articles withdrawn for image issues, 246 (87.5%) were withdrawn between 2015-2020 (2016 is the first year with a significant number of withdrawal notes, almost half of those published by in that year) and the period 2016-2020 was characterized by a slight decrease in problematic articles (74, 26.3%). Elsevier did not have any articles withdrawn in 2020 due to image problems. These data suggest an increased efficiency in dealing with image issues. 2016 seems to mark the beginning of implementing procedures and technologies for image analysis at this publisher(table 7).

**Table 7.** Elsevier image retractions.

|  |  | Retraction Note year |  |  |  |  |  |  |  |  |  |  |  | Total |
| --- | --- | --- | --- | --- | --- | --- | --- | --- | --- | --- | --- | --- | --- | --- |
|  |  | 2010 | 2011 | 2012 | 2013 | 2014 | 2015 | 2016 | 2017 | 2018 | 2019 | 2020 | 2021 |  |
| Publication Year | 2009 | 1 | 1 | 2 | 1 | 1 | 1 | 4 | 3 | 3 | 2 | 2 | 0 | 21 |
|  | 2010 | 1 | 1 | 2 | 2 | 3 | 3 | 16 | 3 | 1 | 1 | 2 | 0 | 35 |
|  | 2011 | 0 | 0 | 0 | 3 | 3 | 4 | 7 | 4 | 4 | 6 | 4 | 0 | 35 |
|  | 2012 | 0 | 0 | 2 | 3 | 1 | 1 | 10 | 7 | 3 | 3 | 5 | 0 | 35 |
|  | 2013 | 0 | 0 | 0 | 2 | 3 | 5 | 2 | 6 | 2 | 2 | 7 | 0 | 29 |
|  | 2014 | 0 | 0 | 0 | 0 | 1 | 3 | 2 | 5 | 5 | 4 | 4 | 0 | 24 |
|  | 2015 | 0 | 0 | 0 | 0 | 0 | 1 | 4 | 4 | 6 | 8 | 5 | 0 | 28 |
|  | 2016 | 0 | 0 | 0 | 0 | 0 | 0 | 2 | 5 | 4 | 10 | 5 | 0 | 26 |
|  | 2017 | 0 | 0 | 0 | 0 | 0 | 0 | 0 | 2 | 6 | 7 | 3 | 1 | 19 |
|  | 2018 | 0 | 0 | 0 | 0 | 0 | 0 | 0 | 0 | 2 | 6 | 8 | 1 | 17 |
|  | 2019 | 0 | 0 | 0 | 0 | 0 | 0 | 0 | 0 | 0 | 1 | 11 | 0 | 12 |
| Total |  | 2 | 2 | 6 | 11 | 12 | 18 | 47 | 39 | 36 | 50 | 56 | 2 | 281 |

In the case of Springer Nature, the focus on images manifested itself a little later (2018-2019), each year from 2009-2020 containing articles withdrawn because of the images(table 8).

**Table 8.**
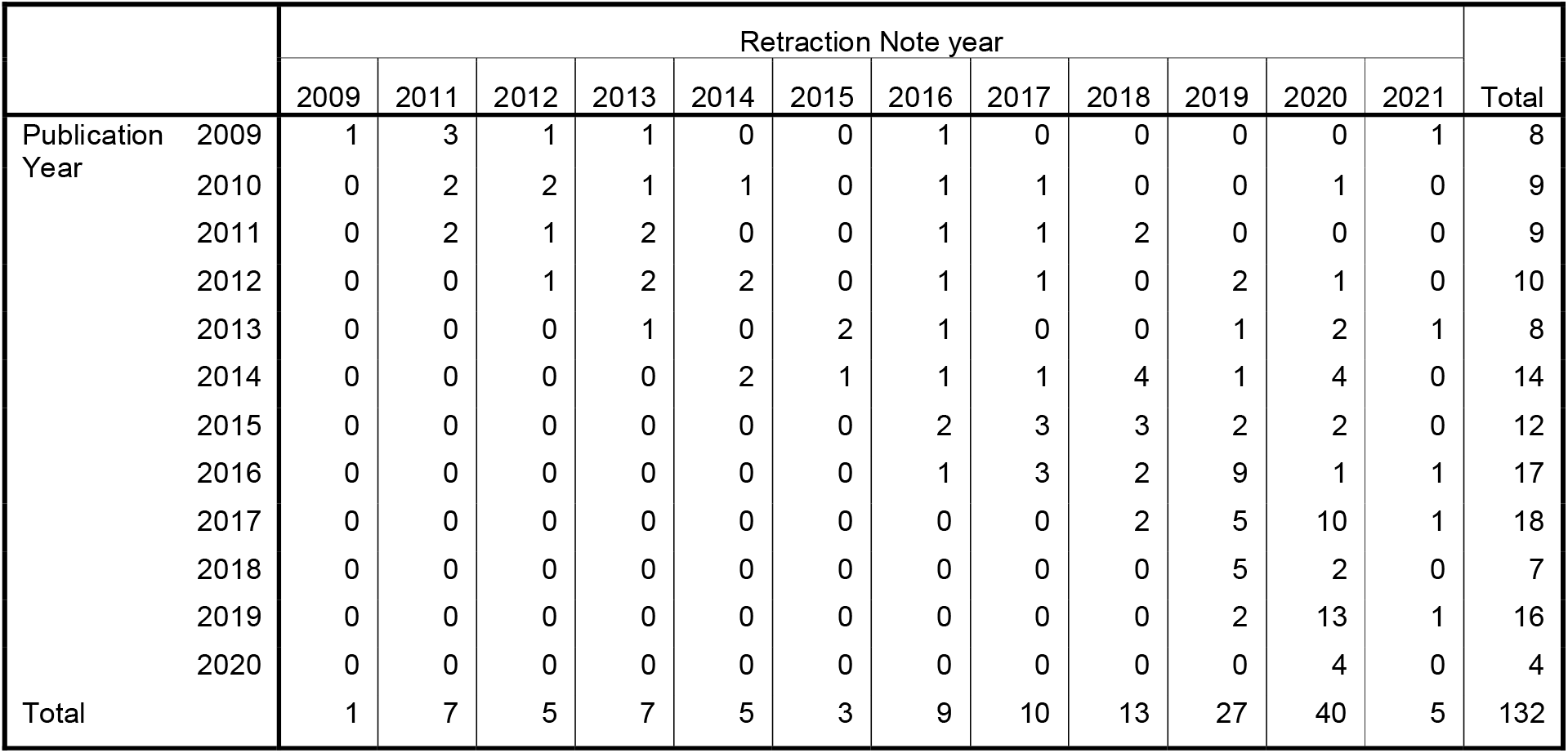
Springer Nature image retractions

**Table 9.**
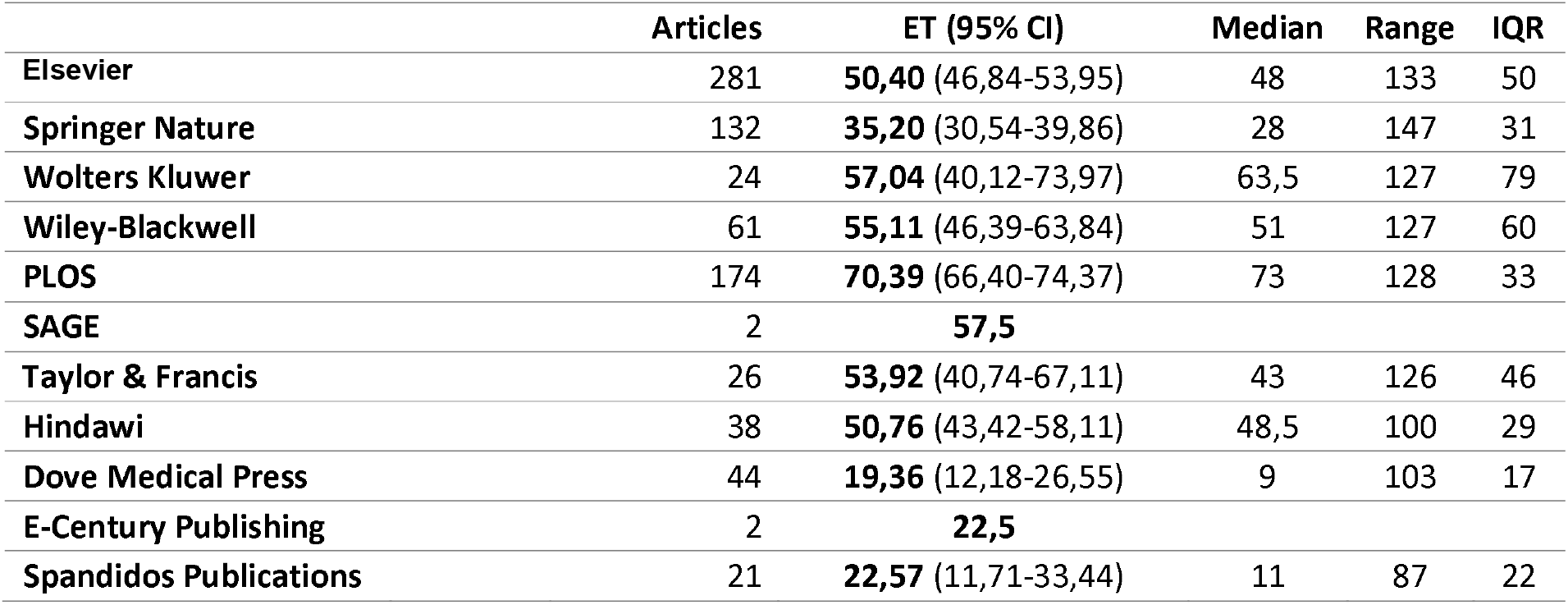
Image retractions by publisher.

Exposure time (ET) for articles with image problems varies between median values of 9/11 months (Dove Medical Press / Spandidos Publications) and 73 months (PLOS). For most other publishers, the median values are between 43 and 63.5 months (average ET value is over 50 months), an exception being Springer Nature, with a median value of 28 months and an average of 35 months.

#### Plagiarism and overlap

The total number of plagiarism and overlap cases for the top 11 publishers is 907 (893 unique articles from which 14 were withdrawn for both plagiarism and overlap): 509 plagiarism, 397 overlap. One of the publishers outperforms the others: with only 22 instances / 21 articles representing less than 10% of their total retracted articles number, PLOS seems to have developed procedures that prevent the publication of articles that reuse text or plagiarize other scientific papers. However, postpublication average exposure time (ET) until withdrawal is the second largest of all publishers: 37.1 months.

Exposure time for plagiarism and overlap cases is presented in table 10.

**Table 10.** Exposure time(ET) for plagiarism and overlap.

|  | Plagiarism | Overlap | Cases | Articles | ET (95% CI) | Median | Range | IQR |
| --- | --- | --- | --- | --- | --- | --- | --- | --- |
| Elsevier | 83 | 92 | 175 | 174 | **26,70** (23,43–29,97) | 20 | 114 | 26 |
| Springer Nature | 138 | 90 | 228 | 222 | **23,34** (20,92–25,75) | 19 | 100 | 24 |
| Wolters Kluwer | 86 | 54 | 140 | 139 | **24,08** (20,04–28,12) | 15 | 105 | 35 |
| Wiley-Blackwell | 30 | 47 | 77 | 77 | **25,17** (19,44–30,9) | 17 | 124 | 26 |
| PLOS | 11 | 11 | 22 | 21 | **37,10** (23,58–50,62) | 37 | 81 | 55 |
| SAGE | 23 | 20 | 43 | 41 | **22,83** (17,52–28,14) | 17 | 65 | 16 |
| Taylor & Francis | 29 | 38 | 67 | 66 | **23,14** (17,58–28,69) | 15,5 | 96 | 28 |
| Hindawi | 47 | 23 | 70 | 69 | **40,14** (33,02–47,27) | 42 | 151 | 50 |
| **Dove Medical Press** | 16 | 18 | 34 | 33 | **17,55** (10,7–24,4) | 10 | 84 | 23 |
| **E-Century Publishing** | 26 | 1 | 27 | 27 | **24,56** (17,04–32,08) | 22 | 63 | 34 |
| **Spandidos Publications** | 20 | 4 | 24 | 24 | **30,13** (20,89–39,36) | 33,5 | 82 | 40 |

The median values of ET are between 10 months(Dove Medical Press) and 42 months(Hindawi) with major publishers relatively well-positioned: 15 months for Wolters Kluwer, 15,5 months for Taylor&Francis, 17 months for Wiley-Blackwell and SAGE, 19 months for Springer Nature and 20 months for Elsevier. Average values of ET spread between 17,5 months(Dove Medical Press) and 40,1 months(Hindawi).

Median values for the top 3 retraction reasons are represented in figure 1.

**Figure 1.**
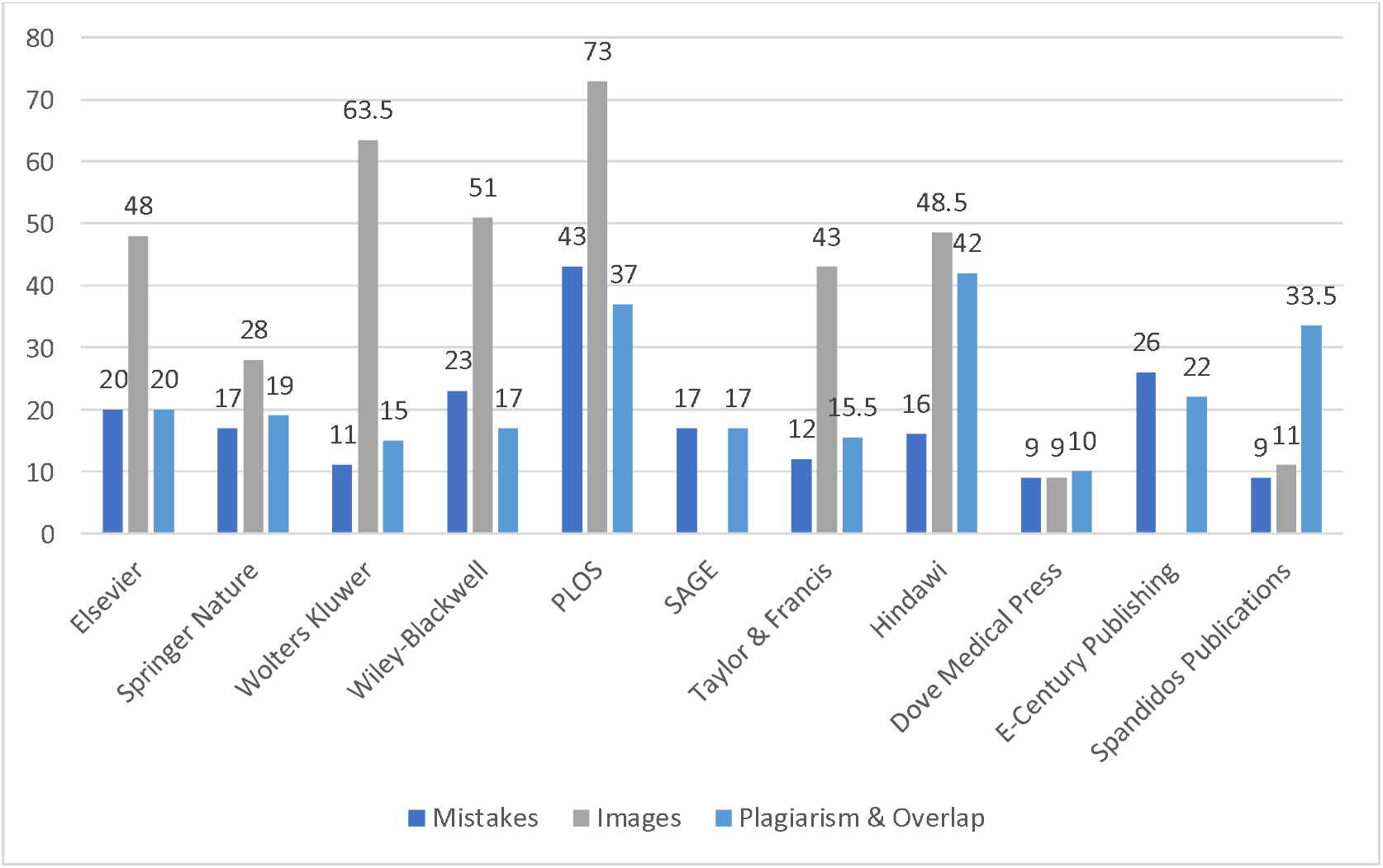
Median values(in months) of top 3 retraction reasons for the top 11 publishers.

### S(peed)D(etection)T(ransparency)P(recision) score

In order to quantify the activity of the publishers, we calculated the SDTP score composed of 6 variables which, in our opinion, can provide an image of their involvement in ensuring the quality of the scientific literature. In order to see the evolution over time, the SDTP score was calculated for the intervals 2009-2018 (3361 articles), 2009-2019 (3931 articles), and 2009-2020 (4844 articles). The score was also calculated for the rest of the publishers below 11th place. The data for 2018, 2019, and 2020 show progress for some indicators and regression of others (see table 11).

**Table 11.**
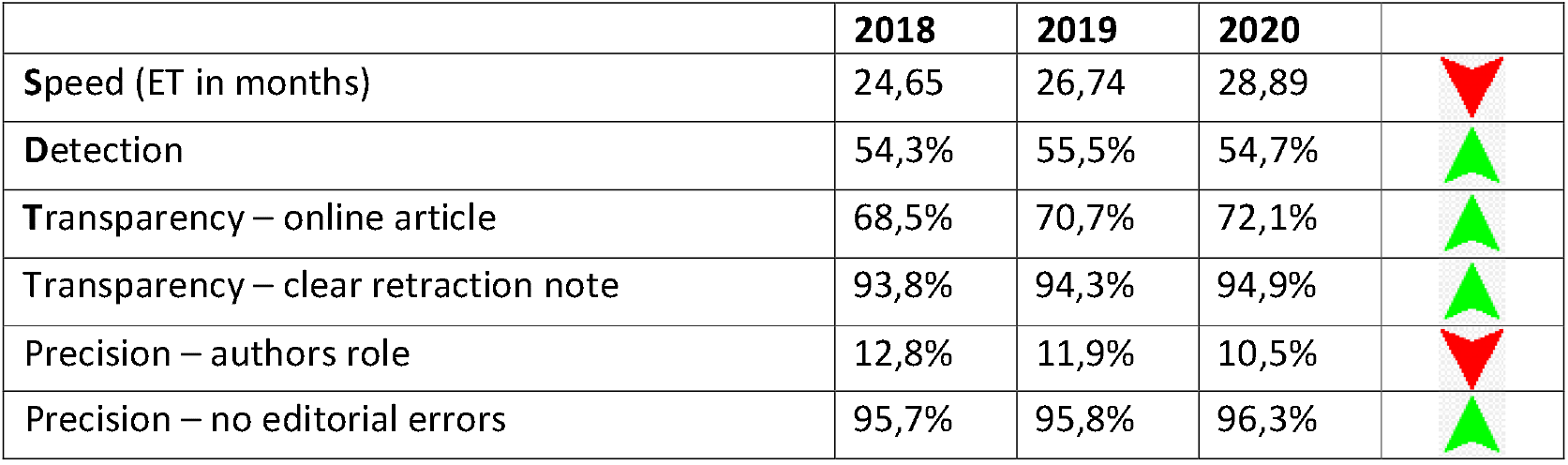
Evolution of main components of SDTP score for 2018-2020.

**Table 12.**
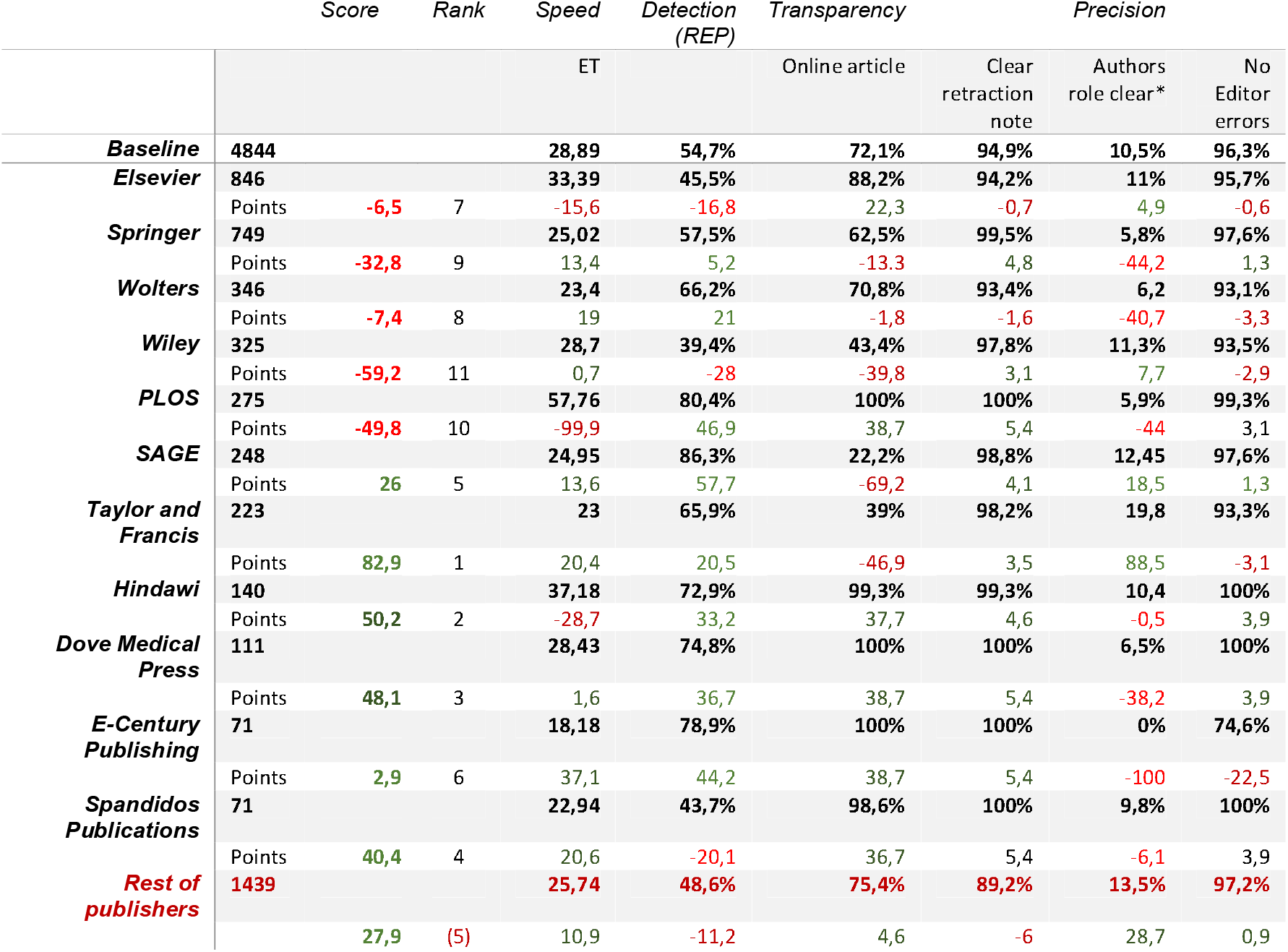
SDTP scores and rank for the period 2009-2020(n=4844). * calculated for 4427 articles with more than one author and no editorial error as retraction reason

**Table 13.**
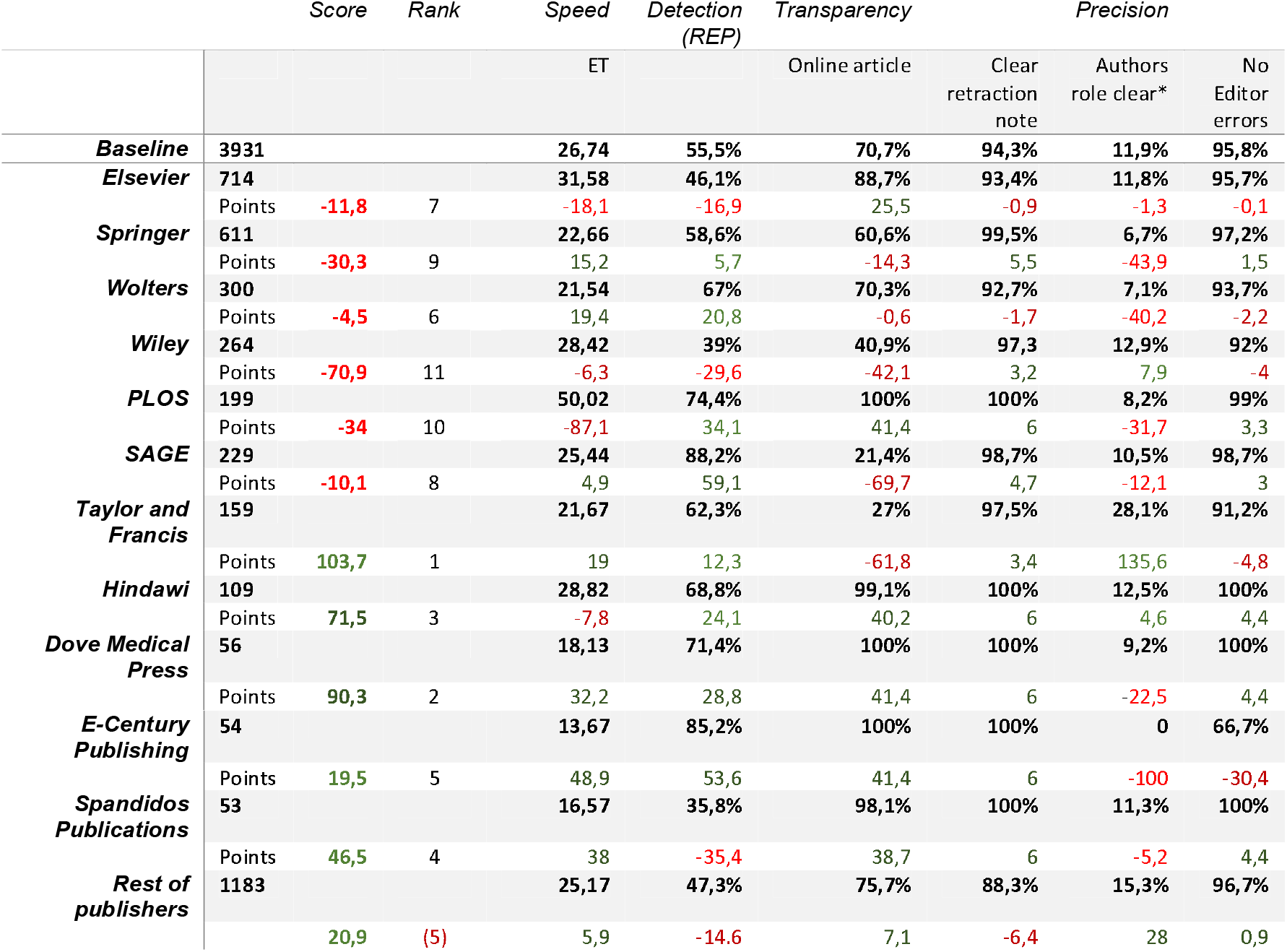
SDTP scores and rank for the period 2009-2019(n=3931). *Calculated for 3569 articles with more than one author and no editorial error as retraction reason.

**Table 14.**
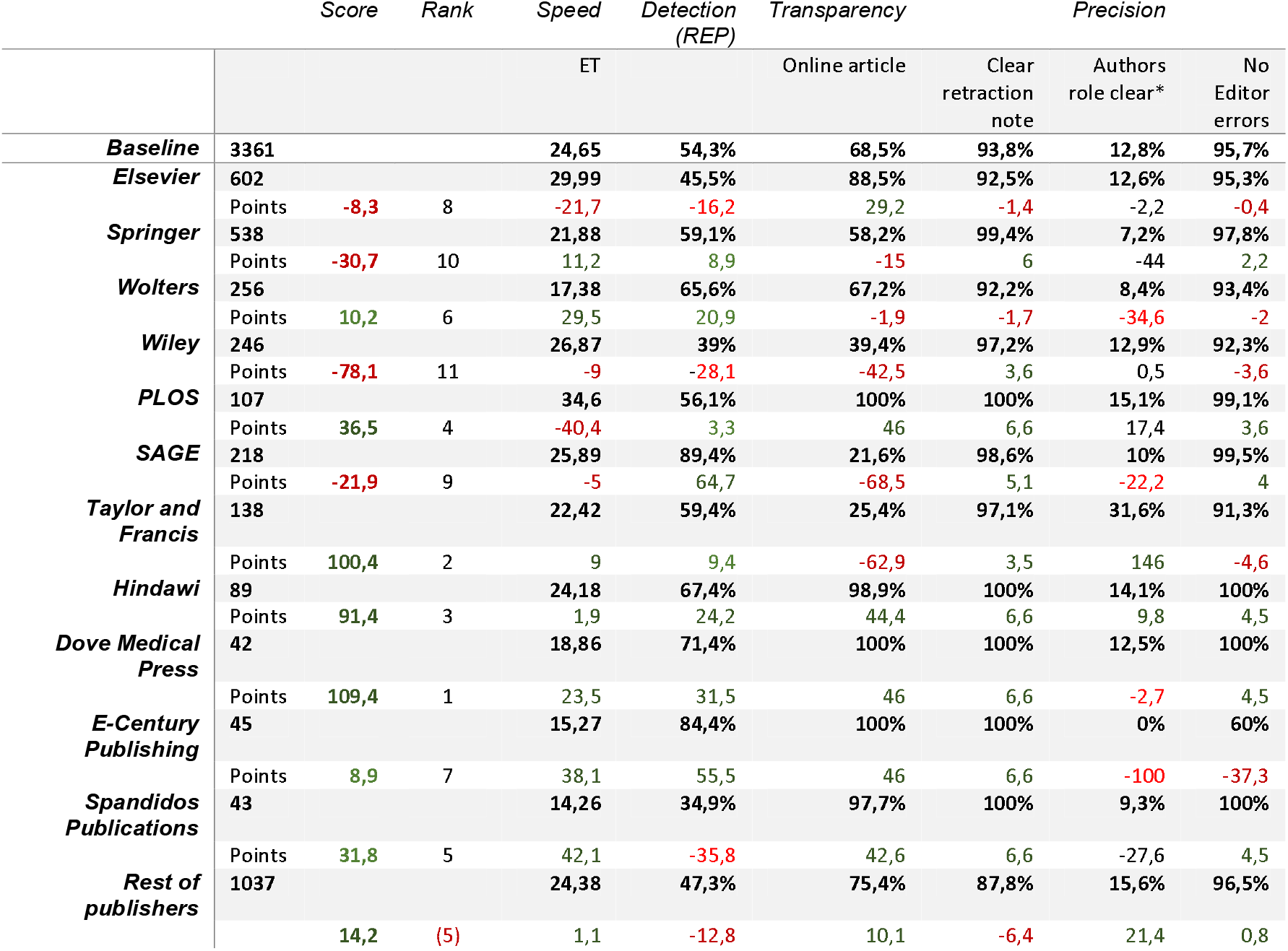
SDTP scores and rank for the period 2009-2018(n=3361). *calculated for 3037 articles with more than one author and no editorial error as retraction reason

The scores for 2018, 2019, and 2020 show signs of a consistent approach (such as Elsevier and Wiley-Blackwell, SAGE, Spandidos Publications), in which the increase in the number of withdrawn items is associated with an improvement in the overall score. There are also signs of a decrease in the quality of the withdrawal notes (such as PLOS, Wolters Kluwer) or the lack of noticeable changes (such as Springer Nature). Some publishers (Taylor & Francis, Hindawi, Dove Medical Press) seem to manage the quality control of their published articles more effectively, recording, however, a decrease of their overall score between 2018 and 2020.

The group represented by the rest of the publishers also marks an increase in SDTP score.

Individual results for the top 11 publishers and the „rest of publishers” group are displayed in tables 15–26(a difference of less than 0,1 points is considered stationary).

**Table 15.** Elsevier 2018-2020 evolution of SDTP score.

|  |  |  |  |
| --- | --- | --- | --- |
| **Elsevier** | **2018** | **2020** |  |
| **Articles** | **602** | **846** | **+40,5%** |
| Speed | -21,7 | -15,6 | 🟢 |
| Detection rate | -16,2 | -16,9 | 🔴 |
| Transparency – online article | 29,2 | 22,3 | 🔴 |
| Transparency – clear retraction note | -1,4 | -0,7 | 🟢 |
| Precision – authors role | -2,2 | 4,9 | 🟢 |
| Precision – no editorial errors | -0,4 | -0,6 | 🔴 |
| **General** | **-8,3** | **-6,5** | 🟢 |
| **Rank** | **8** | **7** |  |

**Table 16.** Springer Nature 2018-2020 evolution of SDTP score.

|  |  |  |  |
| --- | --- | --- | --- |
| **Springer Nature** | **2018** | **2020** |  |
| **Articles** | **538** | **749** | **+39,2%** |
| Speed | 11,2 | 13,4 | 🟢 |
| Detection rate | 8,9 | 5,2 | 🔴 |
| Transparency – online article | -15 | -13,3 | 🟢 |
| Transparency – clear retraction note | 6 | 4,8 | 🔴 |
| Precision – authors role | -44 | -44,2 | 🔴 |
| Precision – no editorial errors | 2,2 | 1,3 | 🔴 |
| **General** | **-30,7** | **-32,8** | 🔴 |
| **Rank** | **10** | **9** |  |

**Table 17.** Wolters Kluwer 2018-2020 evolution of SDTP score.

|  |  |  |  |
| --- | --- | --- | --- |
| **Wolters Kluwer** | **2018** | **2020** |  |
| **Articles** | **256** | **346** | **+35,1%** |
| Speed | 29,5 | 19 | 🟢 |
| Detection rate | 20,9 | 21 | ➡️ |
| Transparency – online article | -1,9 | -1,8 | ➡️ |
| Transparency – clear retraction note | -1,7 | -1,6 | ➡️ |
| Precision – authors role | -34,6 | -40,7 | 🔴 |
| Precision – no editorial errors | -2 | -3,3 | 🔴 |
| **General** | **10,2** | **-7,4** | 🔴 |
| **Rank** | **6** | **8** |  |

**Table 18.**
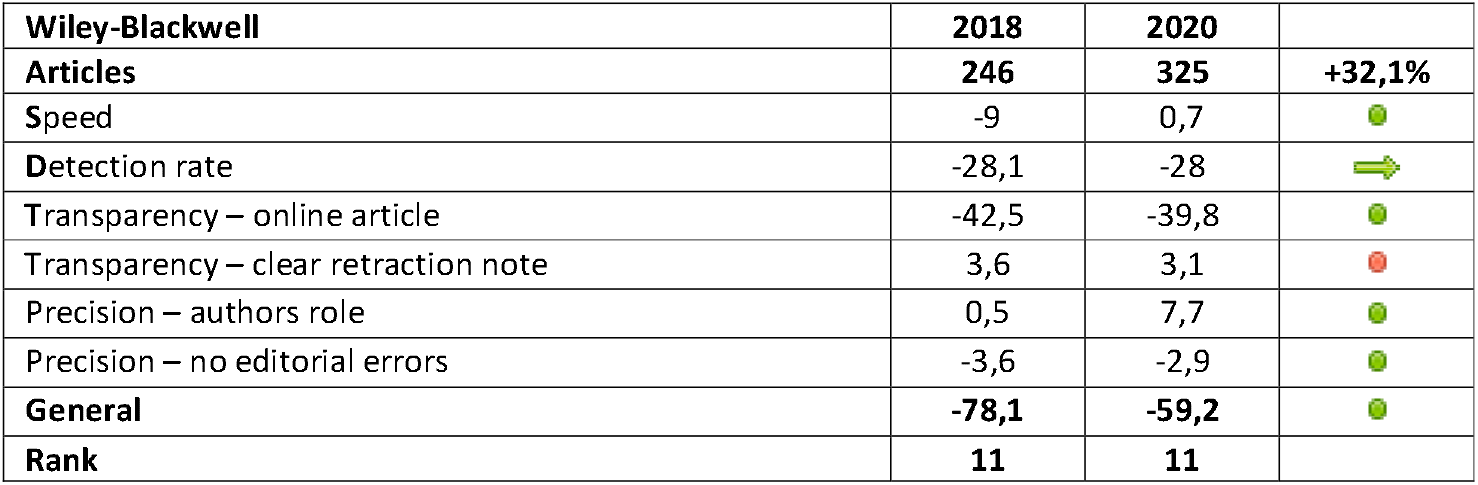
Wiley-Blackwell 2018-2020 evolution of SDTP score.

**Table 19.**
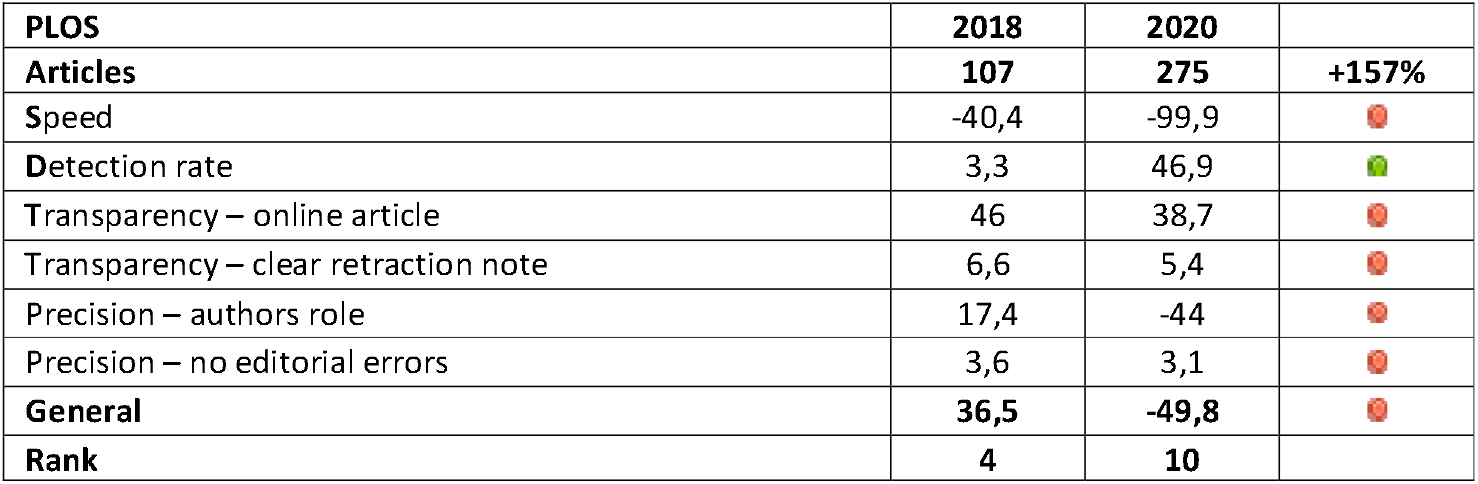
PLOS 2018-2020 evolution of SDTP score.

**Table 20.** SAGE 2018-2020 evolution of SDTP score.

|  |  |  |  |
| --- | --- | --- | --- |
| **SAGE** | **2018** | **2020** |  |
| **Articles** | **218** | **248** | **+13,8%** |
| Speed | -5 | 13,6 |  |
| Detection rate | 64,7 | 57,7 |  |
| Transparency – online article | -68,5 | -69,2 |  |
| Transparency – clear retraction note | 5,1 | 4,1 |  |
| Precision – authors role | -23,2 | -25,4 |  |
| Precision – no editorial errors | -22,2 | 18,5 |  |
| **General** | **-21,9** | **26** |  |
| **Rank** | **9** | **5** |  |

**Table 21.** Taylor&Francis 2018-2020 evolution of SDTP score.

|  |  |  |  |
| --- | --- | --- | --- |
| **Taylor&Francis** | **2018** | **2020** |  |
| **Articles** | **138** | **223** | **+61,6%** |
| Speed | 9 | 20,4 |  |
| Detection rate | 9,4 | 20,5 |  |
| Transparency – online article | -62,9 | -46,9 |  |
| Transparency – clear retraction note | 3,5 | 3,5 |  |
| Precision – authors role | 146 | 88,5 |  |
| Precision – no editorial errors | -4,6 | -3,1 |  |
| **General** | **100,4** | **82,9** |  |
| **Rank** | **2** | **1** |  |

**Table 22.** Hindawi 2018-2020 evolution of SDTP score.

|  |  |  |  |
| --- | --- | --- | --- |
| **Hindawi** | **2018** | **2020** |  |
| **Articles** | **89** | **140** | **+57,3%** |
| Speed | 1,9 | -28,7 |  |
| Detection rate | 24,2 | 33,2 |  |
| Transparency – online article | 44,4 | 37,7 |  |
| Transparency – clear retraction note | 6,6 | 4,6 |  |
| Precision – authors role | 9,8 | -0,5 |  |
| Precision – no editorial errors | 4,5 | 3,9 |  |
| **General** | **91,4** | **50,2** |  |
| **Rank** | **3** | **2** |  |

**Table 23.** Dove Medical Press 2018-2020 evolution of SDTP score.

|  |  |  |  |
| --- | --- | --- | --- |
| **Dove Medical Press** | **2018** | **2020** |  |
| **Articles** | **42** | **111** | **+164,3%** |
| Speed | 23,5 | 1,6 |  |
| Detection rate | 31,5 | 36,7 |  |
| Transparency – online article | 46 | 38,7 |  |
| Transparency – clear retraction note | 6,6 | 5,4 |  |
| Precision – authors role | -2,7 | -38,2 |  |
| Precision – no editorial errors | 4,5 | 3,9 |  |
| **General** | **109,4** | **48,1** |  |
| **Rank** | **1** | **2** |  |

**Table 24.** E-Century Publishing 2018-2020 evolution of SDTP score.

|  |  |  |  |
| --- | --- | --- | --- |
| **E-Century Publishing** | **2018** | **2020** |  |
| **Articles** | **45** | **71** | **+57,8%** |
| Speed | 38,1 | 37,1 |  |
| Detection rate | 55,5 | 44,2 |  |
| Transparency – online article | 46 | 38,7 |  |
| Transparency – clear retraction note | 6,6 | 5,4 |  |
| Precision – authors role | -100 | -100 |  |
| Precision – no editorial errors | -37,3 | -22,5 |  |
| **General** | **8,9** | **2,9** |  |
| **Rank** | **7** | **6** |  |

**Table 25.** Spandidos Publications 2018-2020 evolution of SDTP score.

|  |  |  |  |
| --- | --- | --- | --- |
| **Spandidos Publications** | **2018** | **2020** |  |
| **Articles** | **43** | **71** | **+73,2%** |
| Speed | 42,1 | 20,6 |  |
| Detection rate | -35,8 | -20,1 |  |
| Transparency – online article | 42,6 | 36,7 |  |
| Transparency – clear retraction note | 6,6 | 5,4 |  |
| Precision – authors role | -27,6 | -6,1 |  |
| Precision – no editorial errors | 4,5 | 3,9 |  |
| **General** | **31,8** | **40,4** |  |
| **Rank** | **5** | **4** |  |

**Table 26.**
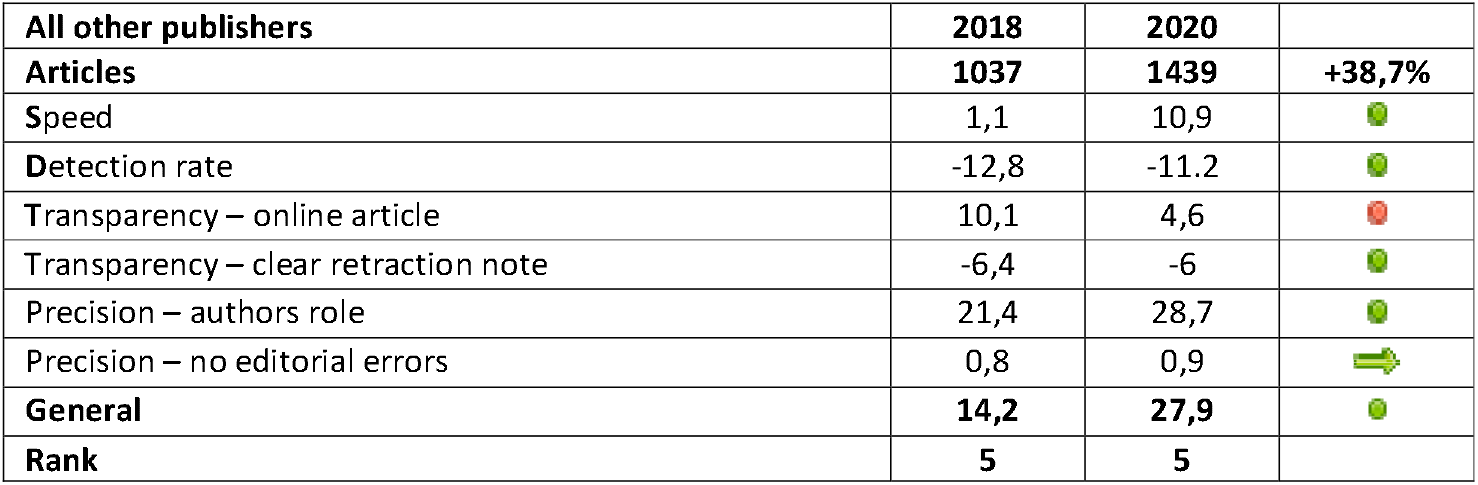
Group of publishers below 11th place, 2018-2020 evolution of SDTP score.

## Discussions

There are many opinions that the actual number of articles that should be withdrawn is much higher than the current number. (Nath et al. 2006; Fanelli 2009; Steen 2011; Poulton 2007; Oransky et al. 2021).

Strengthening editorial procedures may decrease the number of articles withdrawn after publication. The quality and structure of the peer review process (author blinding, use of digital tools, mandatory interaction between reviewers and authors, community involvement in review and registered reports) does have a positive role in preventing the publication of problematic articles. (Horbach and Halffman 2019). However, at the moment, the process of correcting the scientific literature seems to be on an upward trend (Toma and Padureanu 2021) which leads us to believe that there are still enough articles already published that require further analysis.

The post-publication analysis of scientific articles requires considerable effort from publishers/editors. Their performance when it comes to controlling the quality of the scientific product depends not only on internal (organizational) factors but also on external factors such as “Post-publication peer review” or the intervention of authors/institutions(Knoepfler 2015; Teixeira da Silva and Dobránszki 2015). This dependence of quality control on external factors is also reflected in our data by the low involvement rate of the institutions to which the authors are affiliated: out of a total of 4844 withdrawal notes, only 465 (9.6%) mention involvement of an institution. This number may be underestimated, but even so, given that editorial procedures sometimes include communication with authors’ institutions, the lack of effective communication with them makes the work of editors/publishers difficult when it comes to quick withdrawal of an article or clarifying the retraction reasons.

However, despite the impediments generated by the complexity of editorial procedures, if, as suggested (Vuong 2020), withdrawing an article may be regarded as a practical way to correct a human error, it would be probably helpful to measure publisher performance when it comes to quality control of scientific articles.

### How many journals?

We note in our study a concentration of retracted articles in a relatively small number of publishers, the first 11 having 70.3% of the total withdrawn articles and 65.9% of the total number of scientific journals in which these are published. The top 45 publishers account for 88% of all articles and 79% of all journals. In our study, the total number of journals that have retracted at least one article is 1767, representing a small share of the 34768 journals indexed in PubMed (NLM 2022) (https://www.ncbi.nlm.nih.gov/nlmcatalog/?term=nlmcatalog+pubmed[subset]).

Our study included only articles related to human health, excluding 775 articles that did not meet the inclusion criteria. Even if we added another 775 journals (assuming an article/journal), the share would remain extremely low, below 10%. On January 9th, 2022, using the term “Retracted Publication [PT]” we get a total of 10308 records starting with 1951. Using the same logic (one retracted article/ journal) would result, in the most optimistic scenario, another 5464 journals with withdrawn articles, which, added to 1767 in our batch would bring the total to 7,231, just over 20% of the total number of journals registered in PubMed. However, we are helped here by a study published in 2021 (Bhatt 2021), which analyzed 6936 PubMed retracted articles (up to August 2019) and identified 2102 different journals, of which 54,4% had only one article withdrawn.

Our data set contains from September 2019-January 2021 169 journals that retracted at least one article (within the included articles set) and 59 journals with at least one retracted study (within the excluded articles set). Taking into account these figures and including journal overlaps, we can say that the number of journals in PubMed that reported at least one retraction is at most 2330, respectively no more than 6.7% of the total number of active or inactive PubMed journals (the inactive status of a journal is not relevant for our estimation as the evaluation of the number of journal titles took into account the period 1951-2020, the time interval in which all journals had periods of activity).

Mergers, acquisitions, name changes, or discontinuations make it challenging to assess the number/percentage of journals affected by withdrawals from a publishers’ portfolio. In the case of those with a smaller number of titles (PLOS, Spandidos, or Verducci), it is easy to realize that quality problems affect all or almost all of their journals. When we talk about publishers with a medium number (> 100) and a large number (> 1000) of titles, the size of the quality problems of the published articles seems smaller. We are not sure if the data we found for an average publisher (out of 105 journals in the website portfolio (Karger Publishers 2022) or 331 registered journals in PubMed or 94 active journal titles in Scopus, only 8 withdrew a total of 12 articles) or large (out of over 2000 of titles in PubMed only a little over 300 had articles withdrawn) reflect an effective quality control before publication or an insufficient quality control after publication.

Are over 90% of journals without a retracted article perfect? It is a question that is quite difficult to answer at this time, but we believe that the opinion that, in reality, there are many more articles that should be retracted (Oransky et al. 2021) is justified and covered by the actual figures.

### Retraction reasons

Of the 11 publishers analyzed, 9 recorded the highest number of withdrawal notes in 2020, which seems to reflect a growing interest in correcting the scientific literature and indicates the need for a follow-up study to see if new data confirm this trend.

#### Mistakes/Inconsistent Data

Detection of design and execution errors in research (Makin and Orban de Xivry 2019) may stop publishing the article and cause it to be rejected or corrected before publication. However, there are also situations in which the correction is necessary after publication, the errors not being detected in the peer review (Schroter et al. 2008). The leading retraction cause of scientific articles in our study is represented by mistakes / inconsistent data, with 1553 articles. The top 11 publishers have 1005 articles (64.7% of the total).

Data from the top 11 publishers show a large dispersion and an average ET duration of 15.4 months (Taylor & Francis), 17 months (Spandidos Publications), 19.7 months (Wolters Kluwer), and 41.6 months (PLOS). Median values, however, express a better performance for most publishers (table 5). We do not know if the values are due to delays in discovering errors, the length of the withdrawal procedure or a systematic retroactive check implemented at the publisher or journal level, a study on this subject may provide a clearer picture of the correction of errors discovered after publication.

The withdrawal notes mention 229 data fabrication cases, which justifies the need to develop and test the effectiveness of a set of statistical tools capable of detecting anomalies in published data sets (Hartgerink et al. 2019). In our opinion, the number of data fabrication cases may be underestimated: there are 293 cases in which researchers could not provide raw data, 180 cases of lack of reproducibility, lack of IRB approval in 134 cases, 47 cases of research misconduct, and 350 cases of fraudulent peer review. All can camouflage situations where data have been fabricated, even if this was not explicitly mentioned in the retraction. The first 11 publishers have 127/229 (55%) of data fabrication cases.

#### Images

The images represent one of the retraction reasons, which has been growing in recent years, the increased interest in the subject highlighting its unexpected magnitude but also the development of tools that facilitate the detection of image manipulation in scientific articles (Bik et al. 2016; Koppers et al. 2017; Bucci 2018; Bik et al. 2018; Christopher 2018; Sabir et al. 2021).

The total number of image retractions is 1088. Of these, 805 (74%) belong to the top 11 publishers, and 587 of those (54%) belong to 3 publishers: Elsevier (281), PLOS (174), and Springer Nature (132)(see table 9). Exposure time for articles withdrawn due to images is high for all publishers with two exceptions: Dove Medical Press (44 retractions, 19.3 months ET average, median nine months) and Spandidos Publications (21 retractions, ET average 22.6 months, median 11 months). Springer Nature has a slightly better value (132 retractions, mean ET 35.2 months, median 28 months). The other publishers have values between 50 and 57 months, with medians between 43 and 63 months. PLOS has the highest ET: 70 months (median 73).

We reported in a previous study that 83% of image retractions were issued in the period 2016-2020, and the average value of ET is 49.2 months (Toma and Padureanu 2021). Therefore, the values mentioned above are not necessarily surprising, as they are the result of at least two factors: the relatively recent implementation at the editorial/publisher level of image analysis technologies (quite likely started between 2016-2018, see tables 6–8) and the effort of publishers to analyze and withdraw from the literature articles published even 10-11 years ago. The good result of Dove Medical Press can be explained by the fact that most of the withdrawn articles are recently published, and the withdrawal notes were given relatively quickly, in the period 2017-2020, being initiated by the publisher/editor in 33/44 cases. Fast turnaround times appear both at Spandidos Publications and at Springer Nature (table 9). The percentages of initiation of the withdrawal by the publisher/editor are 50% respectively 56%. In these cases, it is possible that there is a workload that is easier to manage, shorter procedures/deadlines, or, simply, a better organization than competitors. The case of PLOS is a special one, the duration of 70 months of ET being 13 months longer than that of the penultimate place (Wolters Kluwer, 57 months); in 114/174 cases, the withdrawal was initiated at the editorial level. The profile of the published articles, the lack of involvement of the authors (only 25/174 cases involved the authors in one way or another), the slowness of the internal procedures, or the too long time given to the authors to correct/provide additional information and data could explain this value. The extended correspondence period with the institutions is not supported by our data (for the 30 cases in which the institutions were involved, the average value of the ET is 72 months).

The values recorded by the other publishers seem to reflect an effort to correct the literature but also difficulties in managing a rapid withdrawal process. Tracking the evolution over time and the factors influencing image retractions ET justifies conducting a follow-up study.

#### Plagiarism and overlap(text and figures, no images)

Detection of plagiarism/overlap can not only be an obligation of the authors/institutions to which they are affiliated (Zimba and Gasparyan 2021) but should also be an essential component of scientific product quality control at the publisher level. The identification of a plagiarized paper / an overlap case after publication represents, in our opinion, a modest editorial performance, especially in the context in which methods and applications (with all their shortcomings) are more and more widespread (Foltýnek et al. 2020; Kulkarni et al. 2021).

The total number of articles withdrawn for plagiarism/overlap is 1201. In 18 of them, both plagiarism and overlap are registered simultaneously. The top 11 publishers have 893 articles (74.3%). In terms of the number of articles, the first place is occupied by Springer Nature (222), followed by Elsevier (174) and Wolters Kluwer (139).

In our study, we identified only one publisher with a good performance in terms of quantity (PLOS - only 8% of all articles withdrawn were plagiarism/overlap). SAGE (17.4% of all withdrawn articles) and Wiley-Blackwell (23.7% of all withdrawn articles) also have a reasonable level.

Regarding the speed of withdrawal of plagiarism/overlap cases, the best performance is recorded by Dove Medical Press (average ET 17.5 months, median ten months). The highest value of ET is recorded in Hindawi (40.1 months on average, median 42 months). Surprisingly, although it has a small number of articles (21), PLOS has the second-highest ET value (37.1 months average, 37 months median). The rest of the publishers have values around the average for the whole lot (average 24 months, median 17 months).

No publisher falls below an average ET of 22 months, with one exception. Their lower performance seems to be influenced, similar to mistakes or image retractions, by the late detection of a small number of articles (skewed distributions with a median lower than average). However, this modest performance shows severe problems at the editorial/publisher level. More than three years to withdraw a plagiarized / overlap article indicates significant gaps in publishers’ detection and intervention capacity in this situation.

### SDTP score

The problem of correcting the scientific literature is one that, by the nature of the procedures to be followed, the resources to be allocated, and the complexity of the interactions needed to withdraw an article sometimes exceeds the organizational capacity of a scientific journal (Wlodawer et al. 2018; Marks 2019; Pelosi 2019; Cooper and Dwyer 2021). We believe that publishers and editors’ early and effective involvement in stopping, discouraging publication, and withdrawing QRP and QPP can help increase the quality of the scientific literature as a whole. The implementation of an independent evaluation system can help such an approach.

The parameters we use to evaluate the performance of publishers aim at the speed of the internal procedures of the journals in their portfolio (speed), the proactive behavior of the editorial staff (detection rate), the transparency, and the precision of the withdrawal notes.

The interval 2018-2020 is characterized by an increase in the number of withdrawn articles by 44%, from 3361 to 4844: the increase varies between 164,3%(Dove Medical Press) and 13,8%(SAGE). At the entire data set level, there is a decrease in performance in ET (an increase from 24.65 months to 28.89 months) and precision (the percentage of identification of responsible authors decreases from 12.8% in 2018 to 10.5% in 2020). The rest of the components are improving (table 11).

For our dataset of PubMed retractions from 2009-2020, the first two places are held by two publishers of different sizes (Taylor & Francis and Hindawi) followed by another medium-sized publisher (Dove Medical Press). The same publishers appear, in changed order, when analyzing the periods 2009-2018 (Dove Medical Press, Taylor & Francis, Hindawi) and 2009-2019 (Taylor & Francis, Dove Medical Press, Hindawi).

We notice that an increase (between 2018 and 2020) in the volume of withdrawn articles has different effects at the publisher level: Dove Medical Press goes from 1st place to 3rd place with a halving of the SDTP score(table 23), PLOS(table 19) goes from 4th place (2018) in 10th place (2020), and its score changes from a positive to a negative value. Taylor & Francis goes from 2nd place in 2018 to first place in 2020 with a slight decrease in the score(table 21), and Spandidos Publication improves its score (table 25). Operating since the end of 2017 at Taylor & Francis (Taylor & Francis 2017), Dove Medical Press seems to have gained an extra speed by almost doubling the number of retracted articles, perhaps due to access to more resources. PLOS performance is affected by late retractions, the average ET increasing from 34.6 months in 2018 to 57.7 months in 2020.

The best and worst performances for the six parameters of the SDTP score are in tables 27 and 28.

**Table 27.** Best SDTP score performance 2009-2020

|  |  | Points |
| --- | --- | --- |
| ET | E-Century Publishing | 37,1 |
| Detection | SAGE | 57,7 |
| Transparency(online article) | PLOS, Dove Medical Press, E-Century Publishing | 38,7 |
| Transparency(clear retraction notes) | PLOS, Dove Medical Press, E-Century Publishing and Spandidos Publications | 5,4 |
| Precision(authors role) | Taylor&Francis | 88,5 |
| Precision(no editorial errors) | Hindawi, Dove Medical Press, Spandidos Publications | 3,9 |

**Table 28.**
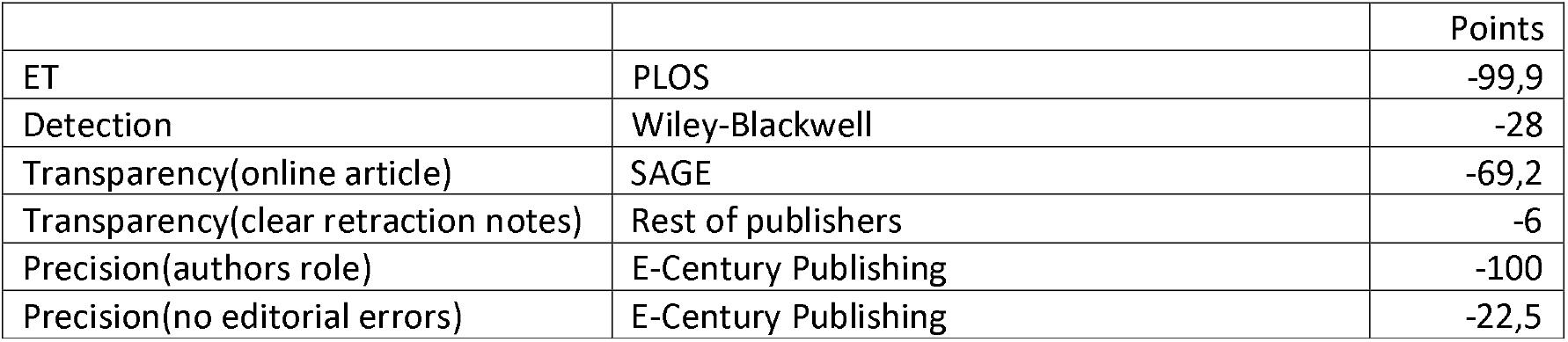
Worst SDTP score performances 2009-2020

|  |  | Points |
| --- | --- | --- |
| ET | PLOS | -99,9 |
| Detection | Wiley-Blackwell | -28 |
| Transparency(online article) | SAGE | -69,2 |
| Transparency(clear retraction notes) | Rest of publishers | -6 |
| Precision(authors role) | E-Century Publishing | -100 |
| Precision(no editorial errors) | E-Century Publishing | -22,5 |

If we look at the individual performance of publishers between 2018 and 2020 (tables 15–26), the picture presented is rather one of declining performance.

Compared to 2018, seven of the 11 publishers have decreased their overall score in 2020. Only four publishers improve their performance between 2018 and 2020: Elsevier, Wiley-Blackwell, SAGE, and Spandidos Publications. The publishers below the 11th place group also show an improvement in the score between 2018 and 2020.

Changes in score components between 2018 and 2020 show interesting developments:

- Only two publishers see an increase in 4 out of 6 indicators (Wiley-Blackwell and Taylor & Francis);
- The other publishers’ group(below 11th place) also has an improvement in 4 indicators;
- Elsevier is the only publisher with three growing indicators
- The other eight publishers register decreases to at least four indicators.

The changes go toward narrowing the gap between 1st place and 11th place: in 2018, the 11th place has −78.1 points, and the 1st place has 109.4 points; in 2020, the 11th place has −59.2 and the first place 82.9 points.

The evolution of scores and indicators, often contrasting with the rank for 2020, associated with the narrowing of the gap between 11th and 1st place, leads us to anticipate a series of developments in the future:

- A possible improvement in the performance of the group of publishers below 11th place;
- A greater homogeneity of results for the first 11 publishers but also for the entire publishing environment;
- An improvement of the results for the big players;
- A continuation of the decrease of specific indicators(like ET) following the appearance of withdrawals for old articles in which the information necessary for a complete withdrawal note can no longer be obtained
- Possible improvement for the involvement of publishers/editors or the editorial errors.

In this context, it is worth discussing whether the time required to withdraw an article reflects the publishers’ performance or there is a need for more complex measuring instruments that consider the multiple dimensions of publishing quality control.

We plan to study these developments in a study that will use retracted articles published between 1.02.2021-31.01.2022 and articles published in the interval 2009-2020 and added in PubMed after 31.01.2021.

## Conclusions

> *„Like a false news report, printed retractions do not automatically erase the error which often pops up in unexpected places in a disconcerting way. There is no instant “delete key” in science.”* (Kiang 1995)

Withdrawal of problematic articles from the scientific literature is a natural process that should involve all stakeholders, including publishers.

Only a small number of journals indexed in PubMed are reporting retracted articles. We estimate that by January 2021, less than 7% of all journals in PubMed had withdrawn at least one article.

However, the correction efforts are obvious for all publishers, regardless of their size.

Exposure time (ET), the involvement of publishers and publishers in initiating withdrawals, the online availability of withdrawn articles, and specifying the responsibility of authors are aspects that can be improved for all publishers reviewed in this paper.

The clarity of the retraction notes and the editorial errors are two indicators for which the potential for progress is limited only to specific publishers.

COPE guidelines must not only be accepted but must also be implemented. In this context, we believe that introducing a reporting standard for retraction notes will allow, along with the introduction of new technologies and the exchange of information between publishers, better quality control of the scientific literature, one that can be easily measured, reproduced, and compared. The SDTP score proposed by us is only a tiny step in this direction.

## Publication bibliography

Ali, Parveen Azam; Watson, Roger (2016): Peer review and the publication process. In Nursing open 3 (4), pp. 193–202. DOI: 10.1002/nop2.51.

Bar-Ilan, Judit; Halevi, Gali (2018): Temporal characteristics of retracted articles. In Scientometrics 116 (3), pp. 1771–1783. DOI: 10.1007/s11192-018-2802-y.

Bhatt, Bhumika (2021): A multi-perspective analysis of retractions in life sciences. In Scientometrics 126 (5), pp. 4039–4054. DOI: 10.1007/s11192-021-03907-0.

Bik, Elisabeth M.; Casadevall, Arturo; Fang, Ferric C. (2016): The Prevalence of Inappropriate Image Duplication in Biomedical Research Publications. In mBio 7 (3). DOI: 10.1128/mBio.00809-16.

Bik, Elisabeth M.; Fang, Ferric C.; Kullas, Amy L.; Davis, Roger J.; Casadevall, Arturo (2018): Analysis and Correction of Inappropriate Image Duplication: the Molecular and Cellular Biology Experience. In Molecular and cellular biology 38 (20). DOI: 10.1128/MCB.00309-18.

Bilbrey, Emma; O’Dell, Natalie; Creamer, Jonathan (2014): A Novel Rubric for Rating the Quality of Retraction Notices. In Publications 2 (1), pp. 14–26. DOI: 10.3390/publications2010014.

Brainard, Jeffrey (2018): Rethinking retractions. In Science (New York, N.Y.) 362 (6413), pp. 390–393. DOI: 10.1126/science.362.6413.390.

Bucci, Enrico M. (2018): Automatic detection of image manipulations in the biomedical literature. In Cell death & disease 9 (3), p. 400. DOI: 10.1038/s41419-018-0430-3.

Christopher, Jana (2018): Systematic fabrication of scientific images revealed. In FEBS letters 592 (18), pp. 3027–3029. DOI: 10.1002/1873-3468.13201.

Cooper, Ashley N.; Dwyer, Jessica E. (2021): Maintaining the integrity of the scientific record: corrections and best practices at The Lancet group. In ESE 47. DOI: 10.3897/ese.2021.e62065.

COPE Council. (2019): COPE Guidelines: Retraction Guidelines. Available online at https://publicationethics.org/files/retraction-guidelines-cope.pdf, checked on December 2021.

Coudert, François-Xavier (2019): Correcting the Scientific Record: Retraction Practices in Chemistry and Materials Science. In Chem. Mater. 31 (10), pp. 3593–3598. DOI: 10.1021/acs.chemmater.9b00897.

Cox, Adam; Craig, Russell; Tourish, Dennis (2018): Retraction statements and research malpractice in economics. In Research Policy 47 (5), pp. 924–935. DOI: 10.1016/j.respol.2018.02.016.

Decullier, Evelyne; Huot, Laure; Samson, Géraldine; Maisonneuve, Hervé (2013): Visibility of retractions: a cross-sectional one-year study. In BMC research notes 6, p. 238. DOI: 10.1186/1756-0500-6-238.

Fanelli, Daniele (2009): How many scientists fabricate and falsify research? A systematic review and meta-analysis of survey data. In PloS one 4 (5), e5738. DOI: 10.1371/journal.pone.0005738.

Fanelli, Daniele (2013): Why growing retractions are (mostly) a good sign. In PLoS medicine 10 (12), e1001563. DOI: 10.1371/journal.pmed.1001563.

Fang, Ferric C.; Steen, R. Grant; Casadevall, Arturo (2012): Misconduct accounts for the majority of retracted scientific publications. In Proceedings of the National Academy of Sciences of the United States of America 109 (42), pp. 17028–17033. DOI: 10.1073/pnas.1212247109.

Foltýnek, Tomáš; Meuschke, Norman; Gipp, Bela (2020): Academic Plagiarism Detection. In ACM Comput. Surv. 52 (6), pp. 1–42. DOI: 10.1145/3345317.

Friedman, Harris L; MacDonald, Douglas A.; Coyne, James C. (2020): Working with psychology journal editors to correct problems in the scientific literature. In Canadian Psychology/Psychologie canadienne 61 (4), pp. 342–348. DOI: 10.1037/cap0000248.

Grieneisen, Michael L.; Zhang, Minghua (2012): A comprehensive survey of retracted articles from the scholarly literature. In PloS one 7 (10), e44118. DOI: 10.1371/journal.pone.0044118.

Hagve, Martin (2020): The money behind academic publishing. Available online at https://tidsskriftet.no/en/2020/08/kronikk/money-behind-academic-publishing, checked on December 27th, 2021.

Horbach, S. P. J. M.; Halffman, W. (2019): The ability of different peer review procedures to flag problematic publications. In Scientometrics 118 (1), pp. 339–373. DOI: 10.1007/s11192-018-2969-2.

International Association of Scientific, Technical and Medical Publishers (2021): STM Global Brief 2021 – Economics & Market Size. Available online at https://www.stm-assoc.org/2021_10_19_STM_Global_Brief_2021_Economics_and_Market_Size.pdf, checked on 11/21/2021.

Karger Publishers (2022): Journals - Current Program. Available online at https://www.karger.com/Journal/IndexListAZ, checked on January 9th, 2022.

Kiang, Nelson Yuan-sheng (1995): How are scientific corrections made? In Science and engineering ethics 1 (4), pp. 347–356. DOI: 10.1007/BF02583252.

Kleinert, Sabine (2009): COPE’s retraction guidelines. In The Lancet 374 (9705), pp. 1876–1877. DOI: 10.1016/S0140-6736(09)62074-2.

Knoepfler, Paul (2015): Reviewing post-publication peer review. In Trends in genetics : TIG 31 (5), pp. 221–223. DOI: 10.1016/j.tig.2015.03.006.

Koppers, Lars; Wormer, Holger; Ickstadt, Katja (2017): Towards a Systematic Screening Tool for Quality Assurance and Semiautomatic Fraud Detection for Images in the Life Sciences. In Science and engineering ethics 23 (4), pp. 1113–1128. DOI: 10.1007/s11948-016-9841-7.

Kulkarni, Sagar; Govilkar, Sharvari; Amin, Dhiraj (2021): Analysis of Plagiarism Detection Tools and Methods. In SSRN Journal. DOI: 10.2139/ssrn.3869091.

Larivière, Vincent; Haustein, Stefanie; Mongeon, Philippe (2015): The Oligopoly of Academic Publishers in the Digital Era. In PloS one 10 (6), e0127502. DOI: 10.1371/journal.pone.0127502.

Madlock-Brown, Charisse R.; Eichmann, David (2015): The (lack of) impact of retraction on citation networks. In Science and engineering ethics 21 (1), pp. 127–137. DOI: 10.1007/s11948-014-9532-1.

Makin, Tamar R.; Orban de Xivry, Jean-Jacques (2019): Ten common statistical mistakes to watch out for when writing or reviewing a manuscript. In eLife 8. DOI: 10.7554/eLife.48175.

Marcus, Adam; Oransky, Ivan (2014): What studies of retractions tell us. In Journal of microbiology & biology education 15 (2), pp. 151–154. DOI: 10.1128/jmbe.v15i2.855.

Marks, David F. (2019): The Hans Eysenck affair: Time to correct the scientific record. In Journal of health psychology 24 (4), pp. 409–420. DOI: 10.1177/1359105318820931.

Mongeon, Philippe; Larivière, Vincent (2016): Costly collaborations: The impact of scientific fraud on co-authors’ careers. In J Assn Inf Sci Tec 67 (3), pp. 535–542. DOI: 10.1002/asi.23421.

Nath, Sara B.; Marcus, Steven C.; Druss, Benjamin G. (2006): Retractions in the research literature: misconduct or mistakes? In The Medical journal of Australia 185 (3), pp. 152–154.

NLM (2022): How do I find a list of journals in PubMed? NLM. Available online at https://support.nlm.nih.gov/knowledgebase/article/KA-04961/en-us, checked on January 9th 2022.

Oransky, Ivan; Fremes, Stephen E.; Kurlansky, Paul; Gaudino, Mario (2021): Retractions in medicine: the tip of the iceberg. In European heart journal 42 (41), pp. 4205–4206. DOI: 10.1093/eurheartj/ehab398.

Pantziarka, Pan; Meheus, Lydie (2019): Journal retractions in oncology: a bibliometric study. In Future oncology (London, England) 15 (31), pp. 3597–3608. DOI: 10.2217/fon-2019-0233.

Pelosi, Anthony J. (2019): Personality and fatal diseases: Revisiting a scientific scandal. In Journal of health psychology 24 (4), pp. 421–439. DOI: 10.1177/1359105318822045.

Poulton, Alison (2007): Mistakes and misconduct in the research literature: retractions just the tip of the iceberg. In The Medical journal of Australia 186 (6), pp. 323–324. DOI: 10.5694/j.1326-5377.2007.tb00917.x.

Rapani, Antonio; Lombardi, Teresa; Berton, Federico; Del Lupo, Veronica; Di Lenarda, Roberto; Stacchi, Claudio (2020): Retracted publications and their citation in dental literature: A systematic review. In Clinical and experimental dental research 6 (4), pp. 383–390. DOI: 10.1002/cre2.292.

Redman, B. K.; Yarandi, H. N.; Merz, J. F. (2008): Empirical developments in retraction. In Journal of medical ethics 34 (11), pp. 807–809. DOI: 10.1136/jme.2007.023069.

Resnik, David B.; Wager, Elizabeth; Kissling, Grace E. (2015): Retraction policies of top scientific journals ranked by impact factor. In Journal of the Medical Library Association : JMLA 103 (3), pp. 136–139. DOI: 10.3163/1536-5050.103.3.006.

Rosenkrantz, Andrew B. (2016): Retracted Publications Within Radiology Journals. In AJR. American journal of roentgenology 206 (2), pp. 231–235. DOI: 10.2214/AJR.15.15163.

Sabir, Ekraam; Nandi, Soumyaroop; AbdAlmageed, Wael; Natarajan, Prem (2021): BioFors: A Large Biomedical Image Forensics Dataset. Available online at http://arxiv.org/pdf/2108.12961v1.

Samp, Jennifer C.; Schumock, Glen T.; Pickard, A. Simon (2012): Retracted publications in the drug literature. In Pharmacotherapy 32 (7), pp. 586–595. DOI: 10.1002/j.1875-9114.2012.01100.x.

Schroter, Sara; Black, Nick; Evans, Stephen; Godlee, Fiona; Osorio, Lyda; Smith, Richard (2008): What errors do peer reviewers detect, and does training improve their ability to detect them? In Journal of the Royal Society of Medicine 101 (10), pp. 507–514. DOI: 10.1258/jrsm.2008.080062.

Shelomi, Matan (2014): Editorial Misconduct—Definition, Cases, and Causes. In Publications 2 (2), pp. 51–60. DOI: 10.3390/publications2020051.

Steen, R. Grant (2011): Retractions in the scientific literature: do authors deliberately commit research fraud? In Journal of medical ethics 37 (2), pp. 113–117. DOI: 10.1136/jme.2010.038125.

Steen, R. Grant; Casadevall, Arturo; Fang, Ferric C. (2013): Why has the number of scientific retractions increased? In PloS one 8 (7), e68397. DOI: 10.1371/journal.pone.0068397.

Teixeira da Silva, J. A. (2016): Silent or Stealth Retractions, the Dangerous Voices of the Unknown, Deleted Literature. In Pub Res Q 32 (1), pp. 44–53. DOI: 10.1007/s12109-015-9439-y.

Teixeira da Silva, J. A.; Dobránszki, Judit (2015): Problems with traditional science publishing and finding a wider niche for post-publication peer review. In Accountability in research 22 (1), pp. 22–40. DOI: 10.1080/08989621.2014.899909.

Teixeira da Silva, J. A.; Dobránszki, Judit (2017): Notices and Policies for Retractions, Expressions of Concern, Errata and Corrigenda: Their Importance, Content, and Context. In Science and engineering ethics 23 (2), pp. 521–554. DOI: 10.1007/s11948-016-9769-y.

Toma, Catalin; Padureanu, Liliana (2021): An Exploratory Analysis of 4844 Withdrawn Articles and Their Retraction Notes. In JSS 09 (11), pp. 415–447. DOI: 10.4236/jss.2021.911028.

Tripathi, Manorama; Sonkar, Sharad Kumar; Kumar, Sunil (2019): A cross sectional study of retraction notices of scholarly journals of science. In DESIDOC Jl. Lib. Info. Technol. 39 (2), pp. 74–81. DOI: 10.14429/djlit.39.2.14000.

van Noorden, Richard (2013): Open access: The true cost of science publishing. In Nature 495 (7442), pp. 426–429. DOI: 10.1038/495426a.

Vuong, Quan-Hoang (2020): Reform retractions to make them more transparent. In Nature 582 (7811), p. 149. DOI: 10.1038/d41586-020-01694-x.

Vuong, Quan-Hoang; La, Viet-Phuong; Ho, Manh-Tung; Vuong, Thu-Trang; Ho, Manh-Toan (2020): Characteristics of retracted articles based on retraction data from online sources through February 2019. In Sci Ed 7 (1), pp. 34–44. DOI: 10.6087/kcse.187.

Wager, Elizabeth; Williams, Peter (2011): Why and how do journals retract articles? An analysis of Medline retractions 1988-2008. In Journal of medical ethics 37 (9), pp. 567–570. DOI: 10.1136/jme.2010.040964.

Williams, Peter; Wager, Elizabeth (2013): Exploring why and how journal editors retract articles: findings from a qualitative study. In Science and engineering ethics 19 (1), pp. 1–11. DOI: 10.1007/s11948-011-9292-0.

Wlodawer, Alexander; Dauter, Zbigniew; Porebski, Przemyslaw J.; Minor, Wladek; Stanfield, Robyn; Jaskolski, Mariusz et al. (2018): Detect, correct, retract: How to manage incorrect structural models. In The FEBS journal 285 (3), pp. 444–466. DOI: 10.1111/febs.14320.

Zimba, Olena; Gasparyan, Armen Yuri (2021): Plagiarism detection and prevention: a primer for researchers. In Reumatologia 59 (3), pp. 132–137. DOI: 10.5114/reum.2021.105974.

